# Exosome-mediated hematopoietic rejuvenation in a humanized mouse model indicate potential for cancer immunotherapy

**DOI:** 10.1101/2020.08.02.233015

**Authors:** Steven J. Greco, Seda Ayer, Khadidiatou Guiro, Garima Sinha, Robert J. Donnelly, Markos El-Far, Sri Harika Pamarthi, Oleta A. Sandiford, Marina Gergues, Lauren S. Sherman, Michael J. Schonning, Jean-Pierre Etchegaray, Nicholas M. Ponzio, Narayanan Ramaswamy, Pranela Rameshwar

## Abstract

Aging is associated with increased morbidity and high economic costs due to a burdened healthcare system and decreased workforce. Parabiotic animal models indicated that secretome from young cells can restore aged tissue functions. We used a heterochronic co-culture system with young and aged mobilized peripheral blood (MPB) or umbilical cord blood (UCB) and showed hematopoietic restoration, independent of allogeneic difference. Bidirectional communication between the aged and young cells influenced the miRNA cargo of exosomes, resulting in partial reprograming of the aged cells. The restored cells enhanced hematopoiesis (e.g., increased lymphoid:myeloid ratio) in immunodeficient mice bearing autologous aged hematopoietic system. Four exosomal miRNAs targeting PAX and PPMIF were partly responsible for the hematopoietic rejuvenation. Notably, increased natural killer (NK) cells within the restored cells eliminated dormant breast cancer cells *in vivo*. The findings could be developed as preventive measure and treatment for sustained immune/hematopoietic competence with potential for immunotherapy.

## Introduction

Aging is a risk factor for chronic diseases, resulting in high morbidity, decreased quality of life and increased health care cost (1). Over time, continuous intracellular stress leads to disrupted tissue physiology such as perturbated tissue homeostasis, stem cell exhaustion, and increased cellular senescence (2). Cumulative age-associated changes could be caused by external and replicative stress that alters the epigenetic dynamics (3). These changes could predispose cells to oncogenic events, bypassing the default protection (4). Regarding treatment for aging, an examination of emerging drugs have underscored the complex process and plausibly demonstrated the challenge of a single drug for cellular rejuvenation (5).

This study focused on methods to rejuvenate the aged hematopoietic system for competent immune system, as a process to improve the overall health. Aging could lead to hematological disorders such as anemia, malignancies, reduced innate immune function and non-hematological disorder, e.g., diabetes and cardiovascular disease (6). An aging hematopoietic system could compromise immune surveillance for emerging malignancies and response to infections (7, 8).

The aged hematopoietic system, partly through defective relationship between hematopoietic stem cells (HSCs) and their niche, leads to loss of HSC quiescence (9, 10). The aging stromal support releases soluble and vesiclular secretome that may contribute to HSC aging (11). The aged have increased HSCs due to loss of quiescence, albeit functionally impaired. Defects of aged HSCs include inefficient serial passaging, loss of heterogeneity, increased genomic mutations, metabolic switch and myeloid bias (2, 10, 12–16). Single driver mutations in aged hematopoietic cells leads to non-malignant clones of indeterminate potential (CHIP), increasing risk for hematological malignancy (17). Other organs can also affect bone marrow (BM) functions. Age-linked dysfunction could occur by the neural system, directly and indirectly by nerve fibers and hormone, respectively (18–20).

The documented defects on hematopoietic aging has not been fully leveraged to reverse and/or halt the aging process (21). Considering the expanded lifespan of humans, fulfilling this gap would have an impact on global public health and the economy. We propose an efficient and non-invasive therapeutic strategy to rejuvenate the hematopoietic system. This method could be used as preventive therapy for middle age individuals and as a potential treatment for the aged population (13).

There is precedence for hematopoietic rejuvenation. Young endothelial cells reversed aged BM endothelial cell-mediated hematopoietic defects (22). Reversed defects of aged HSCs could occur by first dedifferentiating them to iPSCs (23). The recovery of hematopoiesis in young atomic bomb survivors, as well as competence in rejuvenation of aged HSCs were placed in a competent niche underscore the plasticity of HSCs with respect to environmental cues (16, 24). A small molecule against Cdc42 was shown to restore the aging HSCs (25).

We studied hematopoietic rejuvenation with a co-culture method that recapitulated the parabiotic system (26). The latter demonstrated improved cognitive, cardiac and skeletal muscle function of the older animals (27–31). These studies as well as others employing partial cellular reprograming did not examine the hematopoietic/immune systems (32).

We report on young mobilized peripheral blood (MPB) or umbilical cord blood (UCB) rejuvenating aged hematopoietic system, independent of allogeneic differences. The rejuvenating paradigm was corroborated by molecular and functional studies, implicating MYC-targeted genes and decreased p53, caused by specific miRNA within exosomes. Restoration of aging hematopoietic system in immune deficient mice (NSG) with miR or restored cells led to improvement of aging hallmarks, senescence, inflammation and balanced myeloid:lymphoid ratio, similar to young humans (10). Collectively, cellular-mediated hematopoietic rejuvenation demonstrates great promise for immune therapy. More importantly, our rejuvenation method also restored the otherwise low natural killer (NK) activity in the aged cells (33). Strikingly, the restored NK cells eliminated chemotherapy resistant dormant breast cancer stem cells *in vivo*. Thus, the implications for applying this hematopoietic restoration method in immune therapy is proposed and objectively discussed.

## Results

### Hematopoietc restoration of aged MPBs

Clonogenic assays for CFU-GM (granulocytic monocytic) and BFU-E (early erythroid) with MPBs from aged and young donors indicated similarity for CFU-GMs and significantly (*p*<0.05) less BFU-E in young, relative to aged samples (Figures S1A/B). Flow cytometry indicated similar frequency of CD34+/CD38-cells (surrogate of HSCs) but significantly (*p*<0.05) more CD34+/CD38+ (progenitors) in young (Figures S1A/B).

To test aged restoration, we used a transwell co-culturing system with the inner and outer wells containing 10^7^ young and aged MPBs, respectively (heterochronic culture) (Figure 1A). Control isochronic cultures contained aged or young MPBs in both wells. The separating 0.4 µm membranes allowed for secretome exchange but not cell transfer (Bliss et al., 2016) (Figure S1C). CFU-GM was significantly (*p*<0.05) increased in heterochronic cultures up to wk 5 as compared to isochronic cultures (Figure 1B). Aged cells in the heterochronic cultures were significantly (*p*<0.05) more viable as compared to isochronic cultures (Figure 1C).

**Figure 1.**
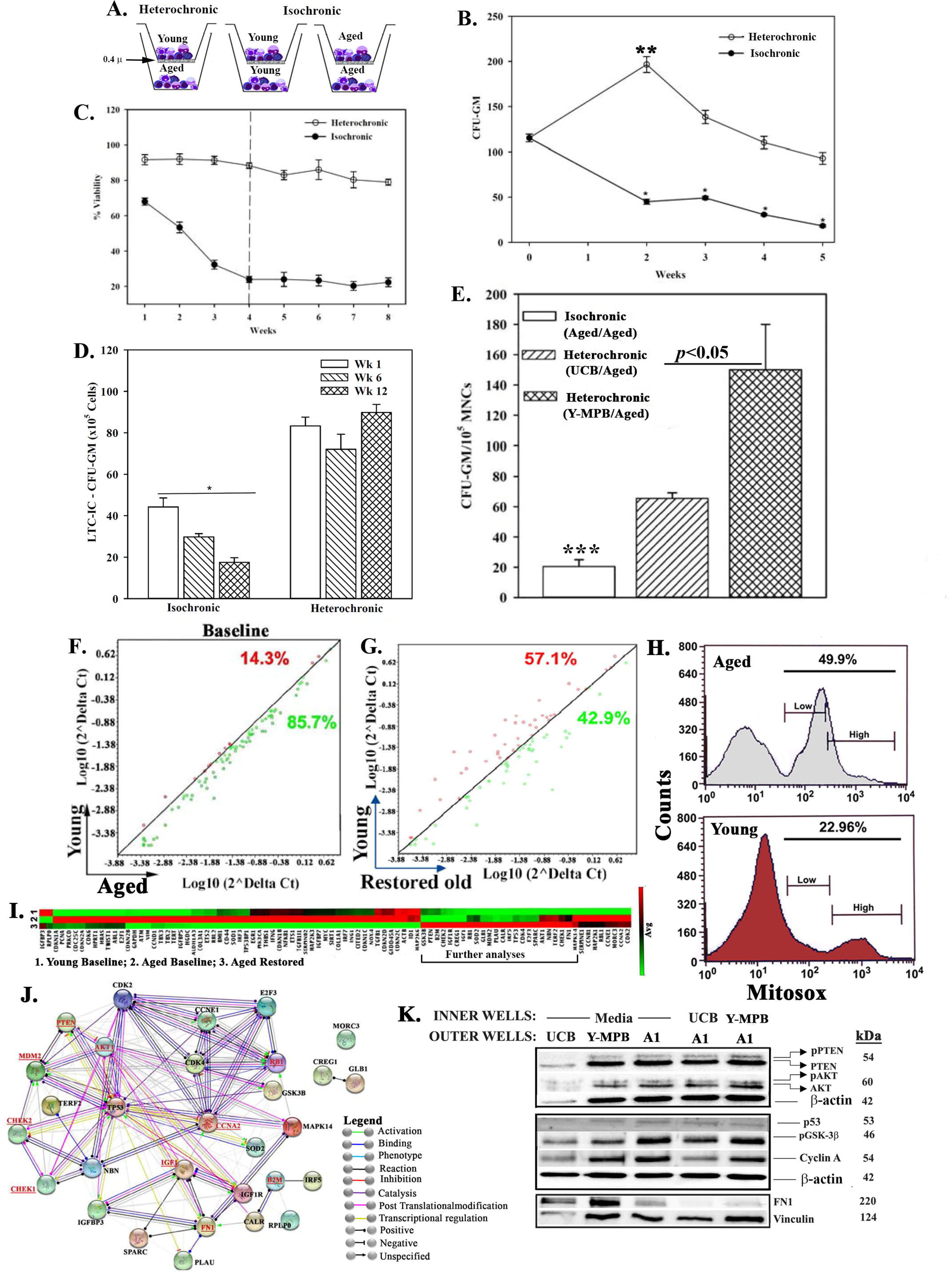
*In vitro* hematopoietic restoration of aged MPBs. (**A**) Cartoon shows the method employed for non-contact isochronic and heterochronic cultures. **(B)** Timeline clonogenic assays for CFU-GM using cells from isochronic (old or young MPBs) and heterochronic cultures (restored old MPBs). The results are presented as mean CFU-GM±SD (n=8 donors, each tested in triplicate/donor; two young donors). **\*** *p*<0.05 vs. similar time points in heterochronic cultures; ** *p*<0.05 vs. time 0 and wks 2, 3 and 4. **(C)** The assay in ‘B’ was repeated except for assessment of cell viability by cell titer blue. The results are presented as mean %±SD (Each time point for a heterochronic and isochronic culture was tested in quadruplicate. **(D)** LTC-IC cultures were established as in ‘A’ except for seeding of the aged MPBs on confluent γ-irradiated BM stromal cells in the outer wells. At wks 1, 6 and 12, aliquots of viable mononuclear cells were analyzed for CFU-GM in clonogenic assay. The values for each time point were plotted together (8 donors, each tested in triplicates, CFU-GM/10^5^ MPBs±SD. \**p*<0.05 vs. heterochronic. (**E**) Heterochronic and isochronic cultures were established as for ‘A’, except for 10^7^ UCB in the inner wells. At wk 4, aliquots of old MPBs were analyzed for CFU-GM and the results presented as mean CFU-GM±SD for 5 different UCB, each tested in duplicate. \*\*\**p*<0.05 vs. heterochronic cultures with UCB. **(F)** Senescence-related gene expression was performed with 84-gene qPCR arrays using cDNA from restored and unrestored (baseline) aged and young MPBs. Gene expression for 4 donors was determined by calculating the ΔC_t_ between gene-of­interest and housekeeping genes and then plotted as Log_10_(2^ΔCt^). Each dot represents the average gene expression for donors. Baseline comparison for unrestored young vs. aged MPBs is shown in red for higher expression in young and green for higher expression in aged. The line *y=x* indicates no change. (**G**) The analyses described in ‘F’ was performed for young and restored and the data are similarly presented. (**H**) Oxidative stress by mitosox assay, delineated as mitosox, negative, low and high by flow cytometry. (**I**) Hierarchical clustering with the array data from ‘F and G’. (**J**) The genes upregulated in the qPCR array in ‘I’ (open boxed region) were analyzed by RAIN, showing predicted interactions. (**K**) Western blot (3 biological replicates) with whole cell extracts from unrestored young MPB (Y), UCB and restored A1 (restored with UCB or Y-MPB). SDS-PAGE: top, 15%; middle 12%; bottom, 6%. See also Table S1, Figures S1 and S2.

We asked if hematopoietic cells (HSCs) were also restored in LTC-IC assay, which is an *in vitro* surrogate method of HSC function (34). The outer transwell was modified for the LTC-IC cultures. At wks 1, 6 and 12, aliquots of cells in the outer wells were assayed for CFU-GM (n=8, each in triplicate). Each time point showed a significant (*p*<0.05) increase in CFU-GM for the heterochronic cultures, relative to isochronic cultures (Figure 1D). Since wk 12 CFU-GMs would be derived from the seeded HSCs, the results indicated restoration of LTC-IC cells, in addition to progenitors.

### Independence of restoration to allogeneic differences of donors

Due to MHC-II expression on HSCs and progenitors, we asked if allogeneic differences between donors could influence restoration via transfer of young MHC-II to stimulate immune cells within the aged cells (35, 36) (Figure S1D). Evidence of MHC-II transfer in one-way MLR with restored aged MPBs as stimulators and unrestored/freshly thawed aged autologous MPBs as responders, showed no response (Figure S1E).

Umbilical cord blood (UCB) HSCs express higher MHC-II, relative to other sources (37), which we corroborated (Figure S1F). We surmised that if allogeneic MHC-II contributed to restoring aged MPBs then UCB should be more efficient as facilitator cells. Heterochronic cultures with UCB cells in the inner wells resulted in increased CFU-GM, but significantly (*p*<0.05) less than young MPBs (Figure 1E), indicating no correlation between MHC-II expression and restoration. One-way MLR with UCB cells showed no evidence of MHC transfer (Figure S1G). MHC-II could be transferred on exosomes (Buschow et al., 2010). However, exosomes from heterochronic cultures showed undetectable MHC-II (Figures S1H/I). Together, the data indicated that allogenicity of the young cells did not contribute to their restorative ability.

### Senescence changes in restored cells

Cellular senescence is an established aging hallmark, which prompted us to analyse restored and unrestored aged MPBs for senescence gene profile (38, 39). An 84-PCR senescence gene array (Figure S2A) indicated 4-fold more expression (∼85%) in the aged vs. young baseline (Figures 1F/S2B). The genes were reduced to 42.9% after restoration (Figure 1G). Using of a 68-antibody array for senescence secretome (SASP) identified 13 baseline proteins in aged MPBs and 4 after restoration (Figures S2D-G) (40) (Figure S2C). Collectively, the restoration paradigm improved the senescence gene profile of aged MPBs.

The vast differences in oxidative stress between young and aged samples led us to further evaluate the senescence PCR array data (Figure 1H). Hierarchical clustering of the averages from biological replicates indicated striking similarities between young and restored aged MPBs with clustering of 30 expressed genes (Figure 1I/cluster in boxed region). We subjected the 30 genes to RNA-protein analysis (RAIN) and found a network consistent with hematopoietic regulatory programs (Figures 1J/S2H). Upregulated MDM2, expected to decrease p53 to avoid cellular senescence without compromise to DNA repair since CHEK1/2 were increased (41). We validated key proteins (Underline, Figure 1J) by Western blot (Figures 1K/S2I). Increases in p-PTEN and p-AKT in restored MPBs suggested a balance proliferation rate. Higher GSK-3β suggested a pathway involving β-catenin to benefit hematopoietic function. Sustained cyclin A suggested no G2 transition, consistent with cycling quiescence in hematopoietic restoration. The link between fibronectin (FN1) and IGF1 was important due to the association of IGF-1 with longevity (42). Since increased FN1 was linked to hematological dysfunction in older individuals, e.g., myelofibrosis, its decrease would benefit hematopoiesis (43).

### RNA in restored cells

In order to gain a better understanding on cell restoration, we performed RNA-seq on biological replicates of restored and unrestored MPBs. Principle component analyses showed distinct clustering within each group (Figure 2A). One young donor with low efficiency as facilitator did not cluster with the other two young donors. Similarity matrix indicated that restored samples moved closer to the young genotype (Figure 2B). Setting a fold change cutoff, described in the methods, and false discovery rate (FDR) <0.05, we identified 2,140 genes, visualized in volcano plots and hierarchical clustering with associated key pathways (Figures 2C/D).

**Figure 2.**
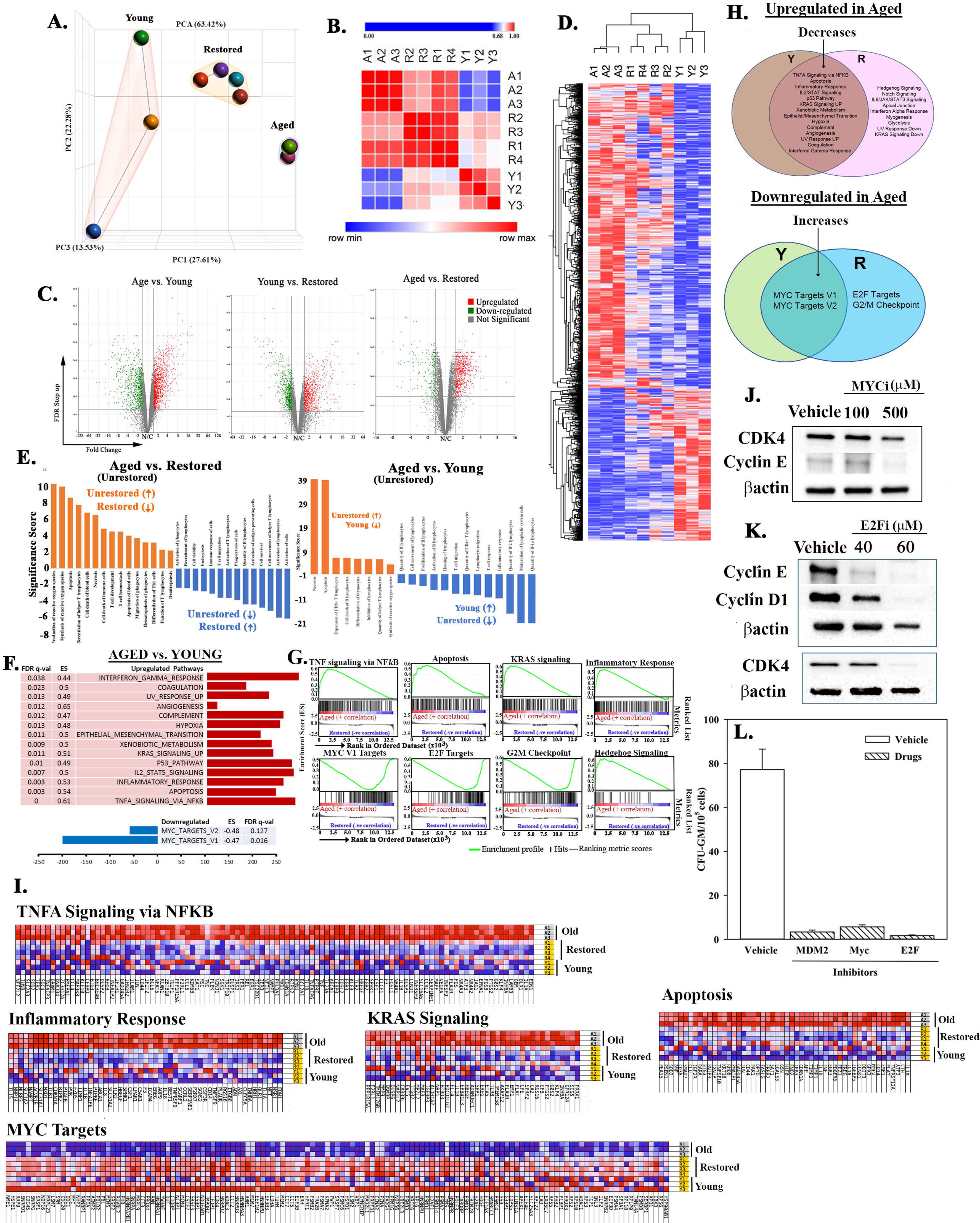
Molecular changes in age-related pathways following restoration. (**A**) PCA of RNA-Seq data from MPB (3 young, 3 age) and 4 restored MPBs. Lines highlight the groups. (**B**) Similarity matrix of ‘A’ for young, age and restored samples. (**C)** Volcano plot of differentially expressed genes. (**D**) Heatmap of fold changes with an FDR ≤0.05 as a cut off with linked significant pathways. (**E**) IPA-determined significant hematological functions with shown comparisons. (**F**) Shown are up- and down-regulated pathways in age, relative to young samples. (**G**) Enrichment plots for the top and down-regulated pathways. (**H**) Venn diagram shows shared and unique pathways between young and restored group, and overlap, changed pathways with restoration. (**I**) Enriched heatmaps of significant changes in ‘G’ (Full heat maps Figure S2). (**J, K**) Western blot for cell cycle proteins with extracts from 3 restored cells, MYC or E2F inhibitors or vehicle. (**L**) Clonogenic assay for CFU-GM with cells restored with 1 µM MDM2, 100 µM MYC or 40 µM E2F, mean±SD (4 different aged donors, restored with 2 young donors; each in triplicates). See also Figure S2.

Ingenuity pathway analyses (IPA) selected the top pathways in hematopoietic development, which confirmed the reversal of age-associated functions such as decreased reactive oxygen species, apoptosis and necrosis (Figure 2E). Gene Set Enrichment Analysis (GSEA) identified significantly up- and downregulated (FDR *q* value <0.05) pathways between aged and young groups. MYC pathways were the only downregulated targets in the aged group but 14 upregulated pathways in the aged vs. young group, with age-associated functions, inflammation, immune suppression and cell death (Figures 2F/G). The five top upregulated pathways were TNFα signaling *via* NFĸB, apoptosis, inflammatory response, IL-2/STAT5 signaling and p53. Restoration was able to correct the age-associated pathways, shown in a Venn diagram of shared and unique pathways (Figure 2H). The original up- and downregulated pathways in the aged samples transitioned into the overlapping section. Heatmaps of the top shared pathways are presented for the enriched sections (Figure 2I, Full heat maps, Figure S2J).

MYC, which was upregulated in the restored cells, could regulate genes linked to partial reprogramming and E2F, regulates cell cycle. We validated their roles in restoration, in the presence or absence of MYC or E2F inhibitors. We included MDM2 inhibitors due to increased MDM2 and decreased p53 in restored samples (Figures 1I/J, 2F). Western blot indicated decreases in proteins linked to G1 transition with MYC and E2F inhibitors (Figure 2J/K). CFU-GM was significantly (*p*<0.05) decrease with the inhibitors, relative to vehicle (Figure 2L). Together, we deduced that hematopoietic restoration was associated with decreased p53 and, increases in MYC and E2F-associated genes.

### Hematopoietic restoration in huNSG

We asked recapitulated the *in vitro* restoration *in vivo* by humanizing NSG mice with CD34+ cells from aged MPB (aged huNSG) (Figures S3A/B) (44, 45). Mice with stable chimerism (19 wks) were transplanted with autologous restored CD3-depleted MPBs (Figures 3A, S3C). There was no untoward pathology, based on survival, body and spleen weights, and pathological examinations of femurs and spleen (Figures S3D-3H).

**Figure 3.**
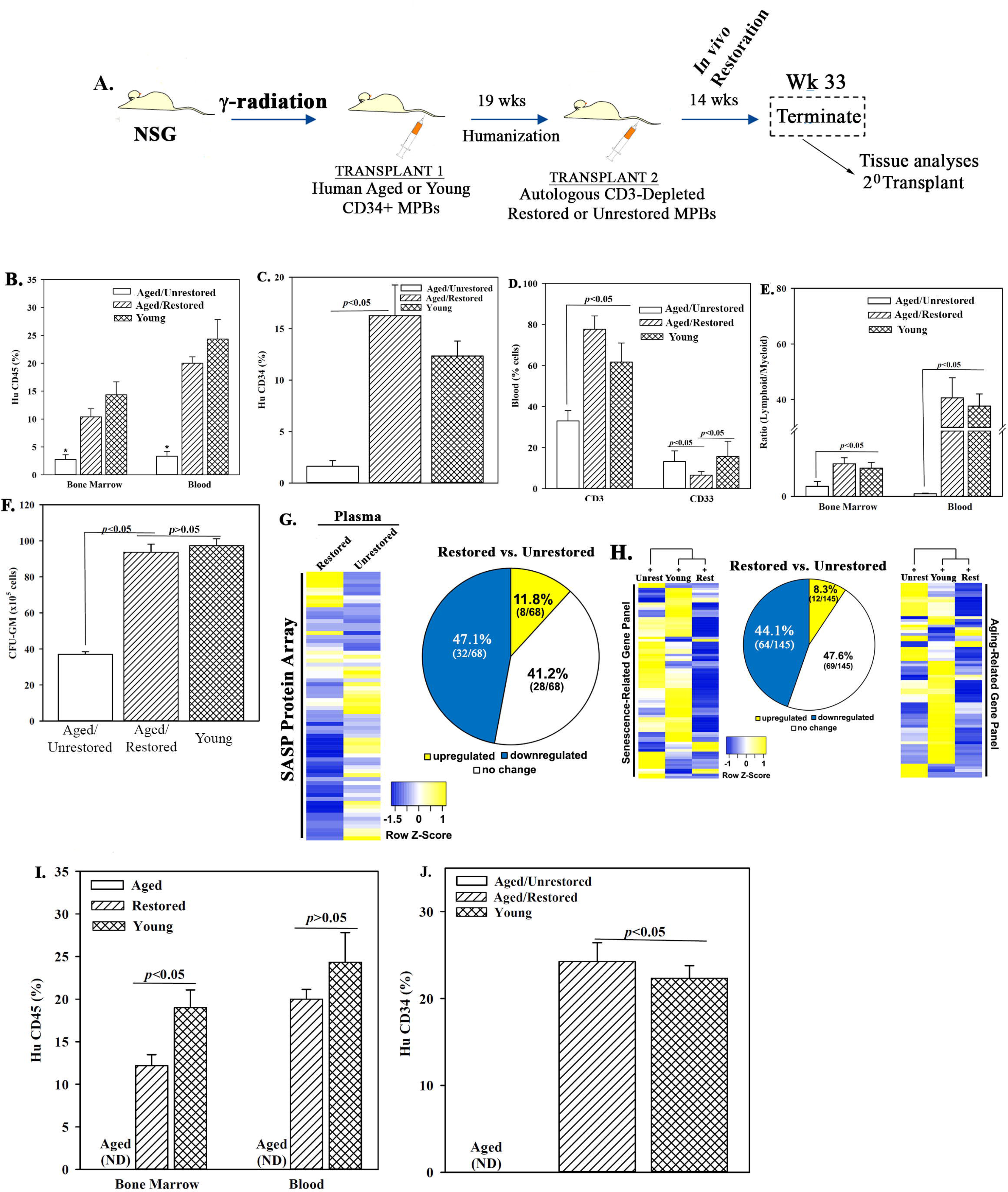
Restored cells transplanted in NSG mice with aged human hematopoietic system (huNSG). (**A**) Overview of the transplantation with aged huCD34+ MPBs then autologous restored (n=12) or unrestored (n=11) CD3-depleted MPBs. Serial transplantation used wk 33 huCD34+ cells. Controls were given young CD34+ cells (n=8). (**B-D**) Flow cytometry for huCD45+ cells in BM and blood, \**p*<0.05 vs. the other groups, (B), huCD34+ cells in BM (C), huCD3+ and huCD33+ cells in blood (D), mean % cells±SD. (**E**) Lymphoid (CD3^+^+ CD19^+^)/myeloid (CD33^+^) ratio in BM and blood. (**F**) CFU-GM in cultures with huGM-CSF and huIL-3 and huCD45+ cells from BM, mean±SD. (**G**) SASF array with huNSG plasma. Semi-quantitative densitometry used 1.5-fold cutoff for classification as up- or down-regulated, or no change. Heatmaps and piechart for differential gene expression. (**H**) RNA from huCD45+ BM cells evaluated qPCR gene arrays. Normalization to housekeeping genes used 1.5-fold cutoff, mean±SEM. \**p*≤0.05 vs. control. (**I & J**) HuCD34+ cells (10^5^), pooled from wk 33 mice were injected into naïve NSG mice (n=3). At 12 wks, mice were analyzed for huCD45+ and huCD34+ cells. ND = non detected. See also Figure S3.

At wk 14, post-second transplant, chimerism was assessed by flow cytometry (Figures S3I-3N). Significant (*p*<0.05) changes in BM and blood of mice with restored cells related to unrestored included: increased huCD45+cells, huCD3 and huCD34+ cells, decreased myeloid cells, and increased lymphoid/myeloid ratio (Figures 3B-F). Clonogenic assays with huCD45+ cells from femurs indicated significantly (*p*<0.05) more CFU-GM in mice with restored cells (Figure 3F). SASP was reduced in the plasma of mice with restored cells (12% in restored vs. unrestored) (Figures 3G, S3O/P). Analyses of qPCR senescence/aging arrays with cDNA from huCD45+ cells of femurs indicated only 8% increased gene expression for mice with restored cells as compared to unrestored cells (Figures 3H, S3P/Q).

We assessed HSC competence by serial transplantation using huCD34+ cells at 33-wk (Figure 3A). Chimerism (based on huCD45 and huCD34+ cells) was achieved in mice given CD34+ cells from primary transplants of young CD34+ cells or aged/restored cells (Figures 3I/J). Human cells were undetectable (ND) with CD34+ cells from primary transplants of aged/unrestored cells (Figures 3I/J). In summary, *in vivo* hematopoietic restoration occurred when mice with an aging hematopoietic system were transplanted with autologous restored MPBs.

### Exosome-containing miR in restoration

Exosomes could migrate through the culture membrane to deliver their cargo, e.g., RNA, proteins and lipids (46, 47). The number and size of exosomes were similar in all cultures (Figures S4A/B). We tested exosomes as mediators of restoration by adding 10^6^ pooled exosomes from day 4 and 7 heterochronic or isochronic cultures to naïve aged MPBs at day 0 and 4. At day 7, CFU-GMs were significantly (*p*<0.05) increased with exosomes from heterochronic and isochronic young cultures, relative to isochronic aged cultures (Figure 4A). CFU-GM was significantly (*p*<0.05) more heterochronic exosomes, relative to young isochronic exosomes.

**Figure 4.**
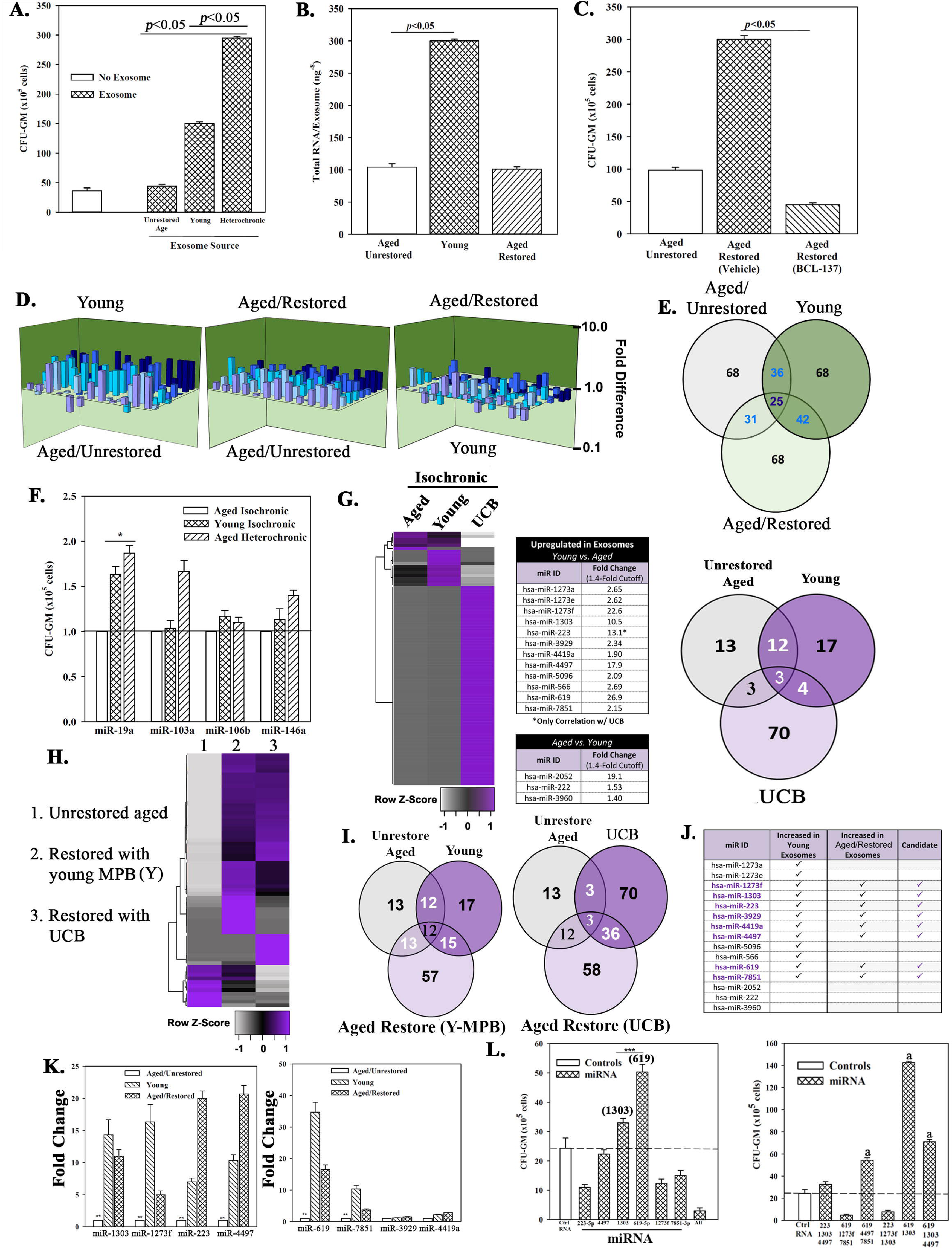
Exosomal miRs in cellular restoration. (**A**) Pooled exosomes (10^6^) from heterochronic or isochronic (young and aged) cultures were added to naïve aged MPBs on day 0 and 4 and at day 7, CFU-GM in clonogenic assays, mean±SD, n=5. (**B**) Total RNA quantification in exosomes from ‘A’, ng^-8^/exosome±SD, n=8. (**C**) BCI-137 or vehicle was added to heterochronic or isochronic cultures, CFU-GM±SD, n=5. (**D**) 3D plots with data from qPCR miR array data using RNA from exosomes. (**E**) Venn diagram showing differential and overlapping miRNAs. (**F**) qPCR for differentially expressed exosomal miRs. Shown are the consistently upregulated miR in young isochronic (dark green bar) and heterochronic cultures (striped bar), mean±SD, n=3. Aged isochronic cultures were assigned a value of 1. \**p*≤0.05 vs. control. (**G**) MiRnome sequencing used small RNA from exosomes of aged and young MPBs or UCB. Heatmaps used miRNA, > 1.4-fold between aged and young samples. Venn diagrams depicts the differential and overlapping miRNAs. (**H, I**) Studies, similar to ‘G**’,** compared miRNAs, sequenced from exosomes of aged isochronic and heterochronic (cultured with young MPB or UCB) samples. (**J**) The 12 miRs showing differential expression between aged and young in ‘G’ were compared to miRs, increased in heterochronic vs. aged isochronic cultures (H, I). Shown are the increased 8 miRs in restored cells. (**K**) qPCR for the 8 miRs, 7 biological replicates, each in triplicate. The data were normalized to miR-7641-2 and presented as fold change using 1 for aged control. (**L**) 6 validated miRs or control miRs were expressed, alone (left) or together (right) in 5 biological replicates, each in triplicate. CFU-GM at day 7, mean±SD. \**p*≤0.05 vs. control. See also Figure S4.

Exosomes from young MPB contained higher RNA content (Figure 4B). We therefore asked if exosomal miRNA could be responsible for hematopoietic restoration (48) by performing heterochronic cultures with BCI-137 (AGO2 inhibitor), which prevents miRNA packaging during exosome biogenesis without affecting their release (Figures S4C/D). At day 7, GFU-GMs were significantly (*p*<0.05) decreased with BCI-137 as compared to vehicle (Figure 4C). The released exosomes contained undetectable miR with BCI-137, pointing to a role for miRNAs (Figures S4E/F).

We sought the causative miRNA(s) with an 84-probe array of commonly expressed miRNAs. Exosomes from isochronic (young and aged) and heterochronic co-cultures showed distinct miR profiles with 25 shared among the groups (Figures 4D/E). IPA predicted targets for the differentially expressed miRs such as p53 pathway (Figure S4G). The latter was consistent with the data shown in Figures 1I-J, 2L. MiR-19a, -103a, -106b and -146a were upregulated in young and restored exosomes from all donors. However, only miR-19a was validated by qPCR (Figure 4F), leading us to expand our analyses with NGS-Seq.

Using an expression cut-off of 100 mappable reads, we identified 13 and 17 unique miRNAs in aged and young exosomes, respectively (Figure 4G). 12/17 exosomal miRs were significantly higher in young than aged, while 3/17 were lower. The miRnome of UCB exosomes were vastly different with 70 miRs in >100 mapped reads but only 4 shared with young MPBs (Figure S4I). Similarly, there were differences in intracellular miRnomes among aged, young, and UCB cultures (Figure S4I). However, upon restoration Young MPBs these baseline differences became shared miR profiles with restoration (Figures 4H, S4J-L).

Among the 12 differentially expressed miRs between young and aged, 8 were up-regulated after restoration (Figures 4G/J). Their validation by qPCR, normalized to miR-7641-2 due to equal expression in all groups (Figure S4M), indicated significant (*p*<0.05) increases in 6 (Figure 4K, annotated in Figure 4J/middle). Among the 6 miRs, only miR-223 and miR-619 were detected in the exosomes of co-cultures in which the restored aged MPBs became the facilitator (Figure S4O).

Cause-effect studies by which ≥1 of the 6 miRs were ectopically expressed in the aged MPBs showed significant (*p*<0.05) increases in CFU-GM with miR-619 and/or miR-1303, and in combination with the other miRs (Figure 4L). MiR-619, -1303 and -4497 became the restorative candidates.

### Targets of restorative miRNAs

We mapped the miR interactome with sequencing data from exosomes and cells (restored and unrestored) and identified top cellular pathways, including T-helper 1 and 2 (Figures 5A, S4P-4S) and, identified molecules that regulate senescence, e.g., CDKN2A and p53. The latter displayed the greatest network convergence between the two datasets (Figure 5B, orange lines), consistent with p53 decrease in the restored cells (Figures 1I/J). Similar molecular outcomes were noted with sequencing data from UCB (Figures S4R-U).

**Figure 5.**
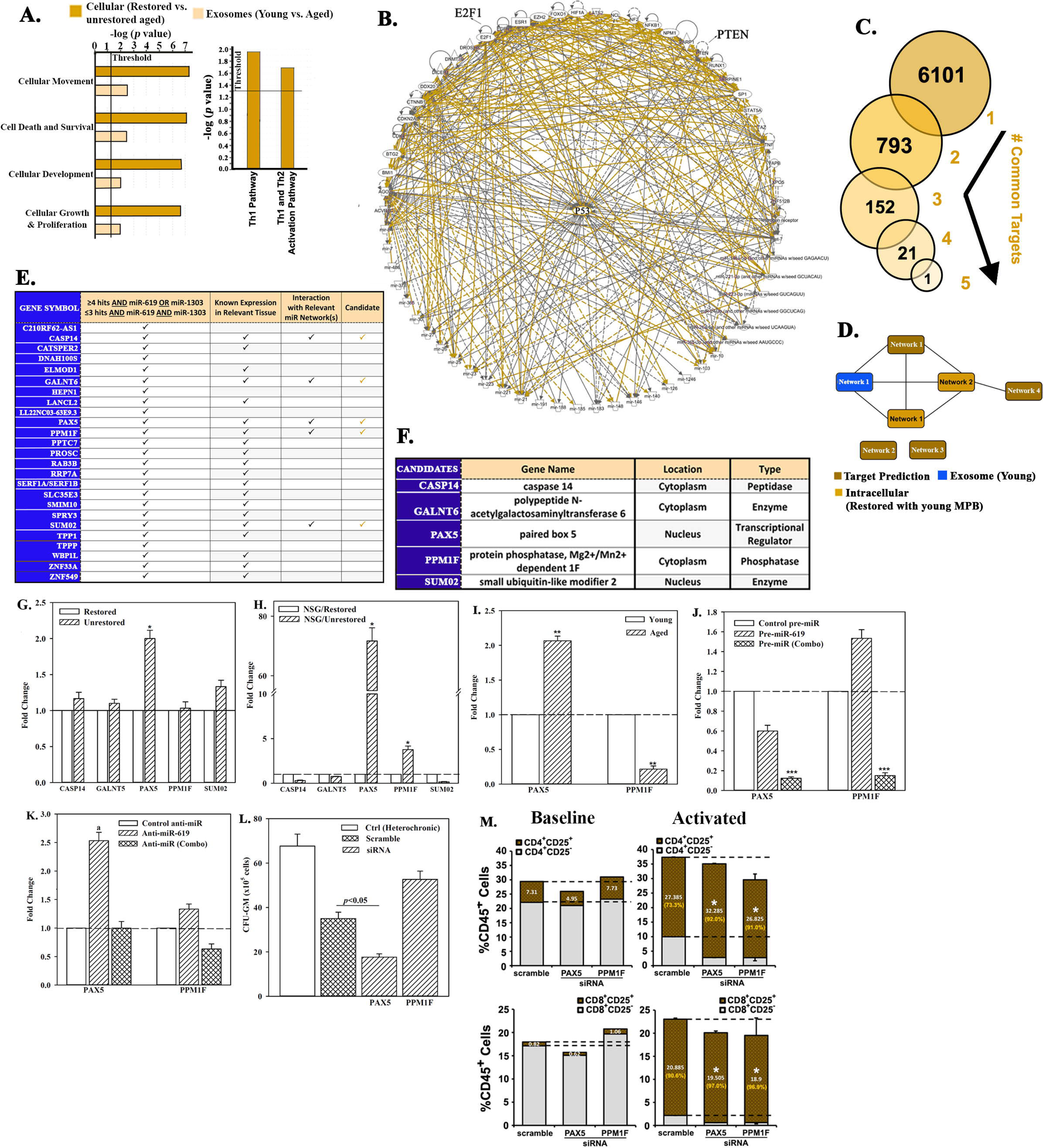
Exosomal miRNA targets in restoration. (**A**) IPA output of top predicted cellular functions (left) and canonical pathways (right) in analyses of exosomal miRNAs from young vs. aged cells, and from cells of heterochronic vs. aged isochronic cultures. (**B**) Radial depiction of young exosomal vs. restored intracellular interactome with p53 at the center of overlapping networks (Orange, direct interactions). (**C**) Analyses of 6 miRNAs (Figure 4L) for targets using TargetScan human database. (**D**) Targets were analyzed by IPA and the predicted networks (brown) compared to the young exosomal (blue) and aged heterochronic intracellular (dark orange) miRNA interactome. (**E**) Tabulation of selected targets and predicted interaction with the miRNA interactome. (**F**) 5 potential downstream targets for functional validation. (**G, H**) qPCR for candidate targets using RNA from aged cells of heterochronic or isochronic cultures (G), or human cells from femurs of huNSG (H), Fold change of normalized (β-actin) results, n=4. (**I**) qPCR for PAX5 and PPM1F in aged and young MPBs, fold change with young donor assigned 1. (**J, K**) Aged MPBs were transfected with pre-miRs or control miR (J) and young MPBs, with anti-miRs or control miR (K). At day 7, the cells were subjected to qPCR for PAX5 and PPM1F, fold change with the control assigned 1. (**L**) Aged MPBs were transfected with PAX5 or PPM1F siRNA or scramble (control). At day 7, the cells were analyzed for CFU-GM. Positive ctrl: heterochronic cultures, mean CFU-GM±SD, n=4. See also Figure S4.

We used target prediction software for direct targets of the 6 restorative miR and identified 6101 targets for individual miR (Figure 5C). We narrowed the targets for common ones then selected those that shared >3 miR hits that included miR-619 and -1303 due to their role in restoration (Figures 5D/4L). The final selection eliminated targets encoding hypothetical proteins and those with restricted expression in neural tissues.

We sought predicted pathways (Figure 5D, brown) in the context of the effector-target interactome (Figures 5D, blue and light orange, 5E). This led to 5 candidate targets (Figure 5F). PCR validation selected the transcription factor PAX5, which was significantly (*p*<0.05) decreased after restoration (Figure 5G). Both PAX5 and PPMIF were decreased in human cells from huNSG BM, with restored cells (Figure 5H). Basal PAX5 and PPMIF in aged and young cells showed an increase of PAX5 and decreased PPM1F in aged, relative to young cells (Figure 5I).

We next asked if the restorative miRs were linked to PAX5 and PPM1F by transfecting aged MPBs with pre-miR-619 or with pre-1303 and -4497 (combo) or, miR mimic (control). MiR-combo significantly (*p*<0.05) decreased PAX5 and PPM1F in the aged MPBs as compared to control or pre-miR-619 alone (Figure 5J). In corollary studies, young MPBs, transfected with anti-miR-619 caused a significant (*p*<0.05) increase PAX5 and PPM1F whereas anti-miR-combo returned to them to baseline (Figure 5K).

Knockdown of PAX5 and PPM1F with siRNA in aged MPBs showed significant (*p*<0.05) decrease in CFU-GM with PAX knockdown and the opposite for PPMIF knockdown (Figure 5L). Since these were aged MPBs with otherwise high level of PAX5, the decrease in the myeloid colonies (CFU-GM) suggested that PAX5, which is involved in early B-cell development might be important at an early progenitor level, hence the decrease in CFU-GM (Auer et al., 2016). However, PAX5 and PPM1F were significant to immune changes in lymphoid cells since their knockdown in aged MPBs led to efficient T-cell activation, suggesting a complex role in hematopoiesis and immune response (Figure 5M). In total, exosomal miRs and their downstream (PAX5 and PPM1F) targets can improve hematopoietic restoration, and their knockdown enhanced lymphoid cell sensitization.

### *In vivo* restoration by candidate miRNAs

Roles for miR-combo (-619 and/or -1303 + -4497) in hematopoietic restoration were tested *in vivo*. Aged huNSG with >1% blood chimerism (huCD45+) were selected for a second transplant with aged MPB, transfected with miR-combo, miR-619 or control miR mimic (Figures 6A, S5A/B). We noted no remarkable pathology in mice (Figures 5SC-H). Compared to control miR, there was significantly (*p*>0.05) higher chimerism (huCD45+) in the BM of mice given miR-transfectants (Figure 6B). MiR-619 transfectants showed significantly (*p*<0.05) higher CD3+ and CD4+ cells and concomitant significant (*p*<0.5) decrease of CD8+ cells, relative to control miR (Figure 6C). The lymphoid:myeloid ratio, which is a hallmark of reversed aging was increased, with a significant (*p*<0.05) increase of CD19+ cells and decreased (*p*<0.05) CD33+ myeloid cells (Figures 6D-F). CFU-GM was significantly (*p*<0.05) increased within huCD45+ cells from femurs of mice given miR-combo-transfected cells or miR-619-transfectants, relative to control miR mimic (Figure 6G). PAX5 level was significantly (*p*<0.05) decrease in mice with miR-619 or miR-combo transfectants (Figure 6H). Similar decrease was observed for PPM1F, but only for miR-combo (Figure 6H).

**Figure 6.**
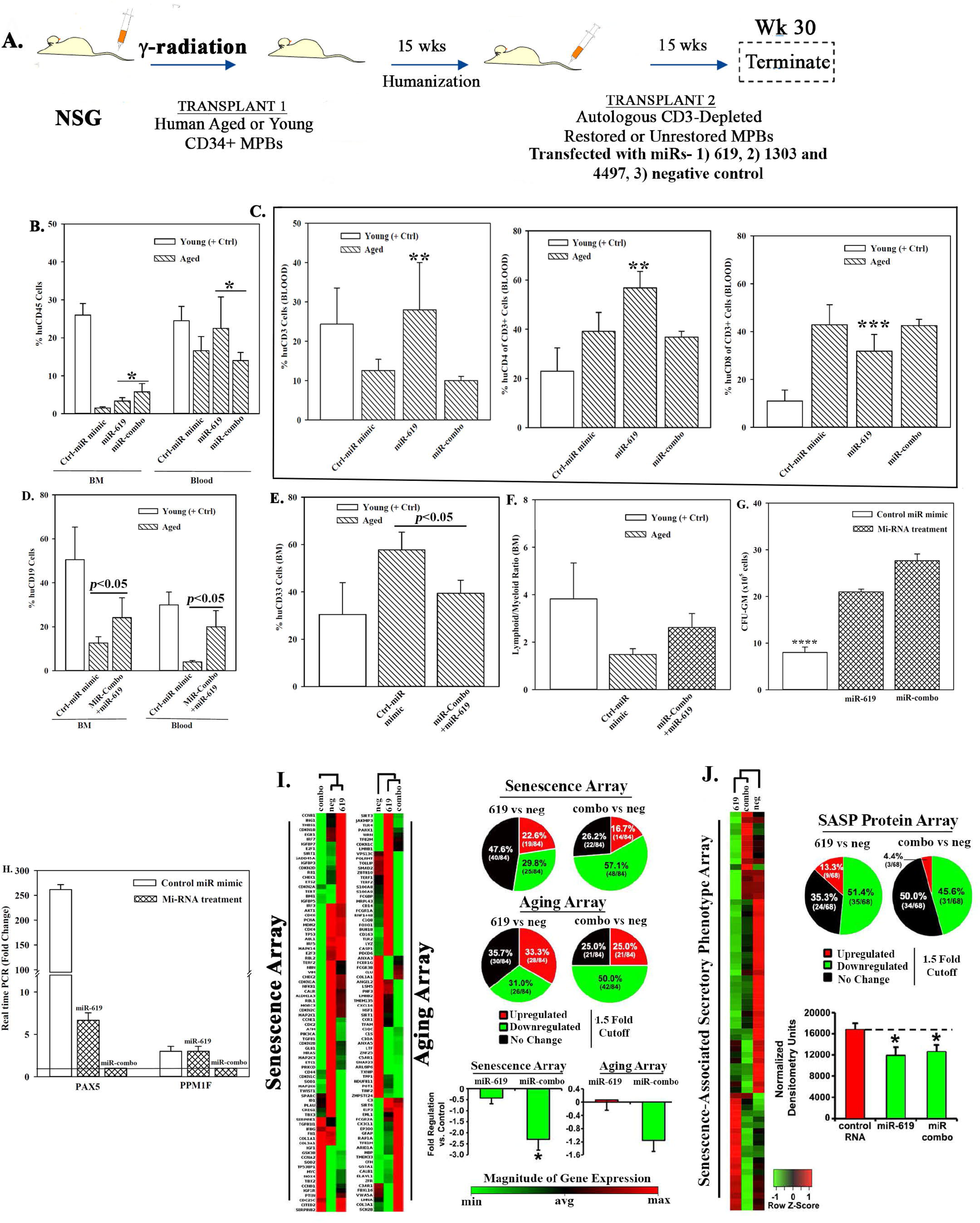
Hematopoietic restoration with miRNAs. (**A**) Protocol with NSG, similar to Figure 3A, also Figure S5. Chimeric mice were given autologous CD3-depleted cells, after 7-day transfection with miR-619, miR-combo, -619, -1303 and -4497, or control (RNA mimic) (n=18). (**B-E**) Mice were analyzed for huCD45 in BM and blood (B); (**C**) huCD3, CD4 and CD8 in blood (C); HuCD19 in blood and BM (D); HuCD33 in BM (E). Results presented as % mean cells±SD (**F**) Lymphoid:Myeloid ratio (CD3^+^+CD19^+^/CD33^+^) in BM. (**G**) CFU-GM with huCD45+ cells from femurs, mean±SD. (**H**) qPCR for PAX5 and PPM1F with RNA from huCD45+cells from femurs. Fold change±SD used 1 for the lowest value. (**I, J**) RNA from ‘H’ were analyzed in qPCR human senescence and aging arrays. Normalized results used 1.5-fold cutoff to classify up- or down-regulation, or no change (I). SASF analyses with plasma. Semi-quantitative densitometry used 1.5-fold cutoff, similar to I (J). Differential gene and protein expressions as heatmaps (left), pie charts (top) and bar graph (bottom), mean±SD. \**p*≤0.05 vs. control. See also Figure S6.

The miR-mediated restoration also reduced cellular senescence, based on PCR array with cDNA from huCD45+ cells of femurs (Figures 6I, S5I-K). MiR-combo led to a significant (*p*<0.05) decrease in >50% of the senescence/age-related genes. Analyses for SASP factors in plasma showed 51.4% and 45.6% decreases with miR-619 and miR-combo, respectively (Figure 6J). Taken together, the miRs restored hematopoiesis, similar to cells from heterochronic co-cultures, and highlighted roles for the transcription factors PAX5 and PPM1F.

### Enhanced Natural killer (NK) activity within restored old MPB in immune therapy

Age-related defects in NK cells can compromise surveillance for emerging malignancy, infection and enhance cell senescence (33, 49). We asked if the restoration process reverted the age-linked NK defects. Flow cytometry indicated 8-fold more CD56+ cells with restoration, relative to unrestored cells (Figure 7A). This correlated with significant (*p* < 0.05) increase in Lytic Units (Figure S6A) in 3/4 aged restored cells (Figure 7B).

**Figure 7.**
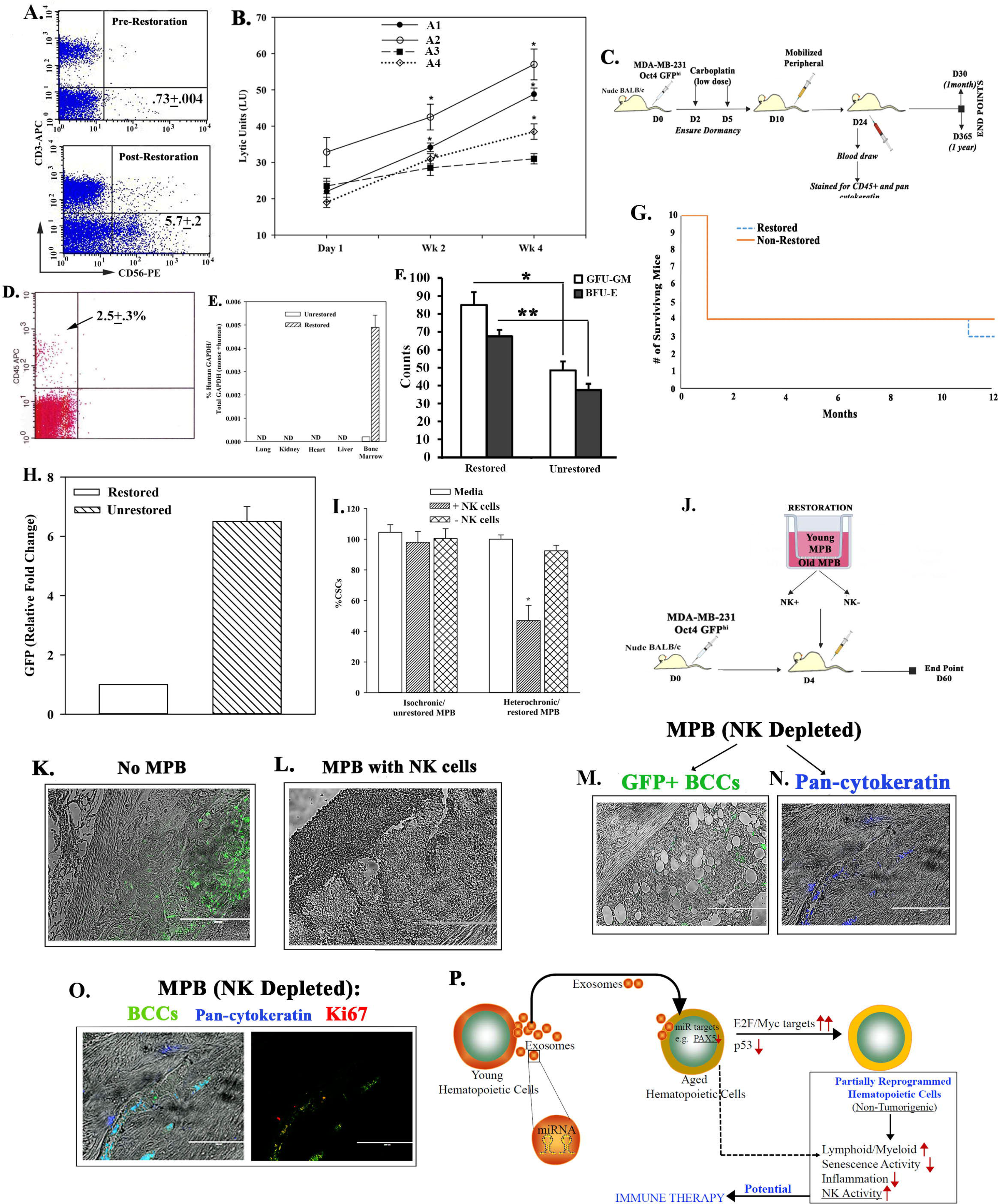
Restoration-mediated targeting of dormant breast CSCs. **(A)** Flow cytometry for CD56+ cells, pre- and post-restoration. (**B**) Timeline LUs (Figure S6A for calculation) in restored cells. **(C)** Treatment protocol with mice harboring dormant CSCs. **(D)** Flow cytometry for huCD45+ cells at 1 month in blood of mice with restored cells. (**E**) qPCR for huGAPDH at 2 months, mean±SD (n=5); ND = not detected. (**F**) CFU-GM and BFU-E in BM at 1 month, mean±SD, n=5. \**p*≤0.05 vs. mice with unrestored cells. (**G**) Survival curve spanning the study period. (**H**) At yr 1, qPCR for GFP with cells from femurs. The results, fold change with the lowest value = 1, mean±SD, 4/group. (**I**) CSCs, co-cultured with restored MPBs (- or + NK cells) for 24 h. % CSCs (GFP+) were determined by microscopy and flow cytometry, mean±SD, n=4. (**J**) Protocol for NSG with dormant CSCs given restored CD3(-) MPB (−/+ NK cells). (**K-M**) GFP (surrogate of CSCs) in paraffin section of femurs, −/+ restored MPB: -MPB (K), +MPB (L), MPB without NK cells (M). (**N & O**) Sections were labeled for pan-cytokeratin-AlexFluor (blue) (N) and Ki67-Rhodamine (O). The latter was merged with bright field image (left). (**P**) Summary: Aged hematopoietic cells instructed young cells to produce specific miRNA containing exosomes to induce MYC and E2F targets for partial reprograming of the aged cells. NK activity is increased. Transplantation of restored cells decreased hallmarks of aging: ↑lymphoid:myeloid, ↓senescence/inflammation. See also Figure S7

We asked if the restored NK cells could target dormant breast cancer (BC) cells (BCCs) in nude BALB/c (50) (Figure 7C). At 1 month, the mice with restored cells were chimeric for human hematopoietic function with no evidence of lethargy, no palpable lymph node and comparable survival between groups (Figures 7D-G). qPCR for GFP in CSCs with pOct4A-GFP indicated 6 fold more GFP+ cells in mice with unrestored MPBs, related to unrestored cells (Figure 7H). Undetectable GFP+ cells in the blood indicated that the unrestored MPBs did not reverse the dormant phase of BC, close to the endosteum (Figures S6B/C). Together, these findings showed the restored MPBs capable of clearing dormant BCCs in BM.

Next, we asked if the cleared BCCs were due to the restored NK cells. At 24 h after co-cultures of CSCs and restored MPBs (NK+ or NK-) (Figure S6D), GFP+ cells were significantly (*p*<0.05) decreased with NK+ MPBs as compared to NK-cells (Figure 7I). Adapting these studies in NSG mice (Figure 7J), we could not detect GFP+ cells in mice given restored NK+ cells whereas control mice (baseline) and mice given NK-restored cells showed dormant CSCs (green) in femurs (Figure 7K-N). The NK-restored cells induced proliferation of the CSCs, based on labeling for Ki67 (Figure 7O). In summary, NK cells in restored MPBs effectively targeted dormant CSCs.

## Discussion

We report on restoration of aged hematopoietic cells with significant improvement of age hallmarks-decreased senescence markers, improved lymphoid:myeloid ratio, decreased inflammation and enhanced NK numbers/activity. We successfully applied the *in vitro* findings *in vivo* in which NSG mice carrying aged human hematopoietic system were functionally improved with transplanted autologous restored cells (Figure 3). The improvement included HSC functions, based on serial passaging of CD34+ cells and, enhanced hematopoietic activity (Figures 3I/J). Despite allogeneic differences between the young and aged donors, restoration was independent of MHC-II variations (Figure 1S). The young cells allowed for cell survival, providing them with an advantage to undergo partial reprograming.

The increased lymphoid cells also included CD19+ B-cells. However, when we compared femurs and spleen and blood, there were relatively less CD19+ cells in the spleen (Figure SK). Since there was prominence in the follicles of spleens, we propose that there might be high activity of B-cell maturation in the spleen for release into the periphery. Although exosomes mediated rejuvenation process, collecting exosomes from young cells is not sufficient since efficient hematopoietic function was noted only when the exosomes were collected from heterochronic cultures, indicated the need for the aged cells to influence the content of the exosomes (Figures 4A/H).

Youthful hematopoietic function is expected to be associated with minimum senescence. However, our ability to reverse the senescence profile of aged hematopoietic cells with young cells was a significant finding since this supported successful restoration. Decreased NK cell number is partly responsible for senescence (33). Questions that remain is whether restoration of NK cells is involved in the youthful changes and whether the senescence cells were eliminated rather than reverting to competence. The latter will be answered in timeline studies to track the aged cells during restoration, including *in vivo* transplantation. Transplantation of restored MPBs within a milieu of an aged human hematopoietic system led to hematopoietic competence that slightly surpassed mice transplanted with young CD34+ cells (Figure 3). There were increased huCD34+ cells in the mice given restored cells as compared to those with unrestored and young cells (Figure 3C). The next two paragraphs expand on plausible mechanisms for enhanced hematopoietic activity in mice given restored cells as a second transplant.

It is possible that exosomes secreted by the transplanted restored cells will restore the aged cells/microenvironment *in vivo*. We based this premise on the ability of exosomes and its miR cargo to rejuvenate hematopoiesis (Figure 4). Interestingly, the restored cells could become facilitator/young cells to restore aged MPBs, indicating the youthful function of the restored cells (Figure S4O). Thus, one can premise that exosomes from the transplanted cells would continue to secrete exosomes to employ *in vivo* restoration of the aged hematopoietic cells (Figures 4-6). If this was the sole method of *in vivo* restoration, then competent HSCs were not expanded during the *in vitro* restoration. Rather, *in vivo* restoration occurred by cellular therapy. Thus, such *in vivo* restoration would be in addition to partial reprograming of the aged cells. Evidence for this is based on functional studies and, molecular data accrued by sequencing. The LTC-IC assay, which is a surrogate of HSC function (34), showed sustained competence of the primitive hematopoietic cells during restoration but similar observation was not noted when the aged MPBs were placed alone in isochronic cultures (Figure 1D).

Sequencing of restored cells identified MYC-linked targets comprise as the only decreased pathways in aged cells, which was reversed with restoration (Figures 2F/G). We were intrigued by this observation as increased MYC-linked pathways correlated with decreased p53 pathways (Figures 1/3). The significance of these two observations is related to the need for decreased p53 and increased MYC to enhance cellular reprograming, which supported partial reprogramming (51) (Figures 1/2). Additionally, pharmacological inhibitors verified roles for MYC and p53 in hematopoietic restoration (Figure 2L). Long-term observation of mice showed no untoward effect such as hematological malignancy. This therefore argues against full reprogramming, similar to induced pluripotent stem cells, which would develop malignancy. The majority of experiments used 7-day co-cultures with results similar to 1-month outcomes, also supporting early partial reprogramming (Figure 1).

Protein and cDNA senescence arrays showed restoration improved cellular senescence and, RNAseq indicated decreases of inflammatory genes, signaling changes in two hallmark groups of aging (52) (Figures 2G/I). More importantly, restoration increased lymphoid:myeloid ratio in the humanized mice (Figure 3). Similar findings were noted when the aged cells were transfected with the identified miRs (Figures 3/6). Increased PU.1/SP-1, which is linked to age-mediated inflammation was decreased with restoration (53).

E2F targets were also increased with restoration (Figure 2). Since E2F activity is silenced during G1 phase of the cell cycle, this suggested the involvement of the cell cycle program during restoration (54). Indeed, there were increases in cycling genes linked to E2F and MYC (Figure 2). MYC could be involved in controlling E2F switch for cyclic control, consistent with what would be expected during restoration (54). In this regard, these findings are intriguing and require robust tracking of the cells during the period when cells transition to partial reprogramming. As the research continues, we will track all cells to fully understand how partial reprogramming contributes to restoration. RNAseq indicated an increase in G2M pathway, which would be consistent with the restored cells undergoing DNA repair. Since p53 was decreased during restoration, we deduced that its role in DNA repair was not compromised due to increases in other repair genes (Figure 1J). The increase in p53 in the aged cells might be linked to apoptosis, noted by associated pathways (Figure 2F/I).

Similarity of CD34+/CD38-progenitors between young and aged MPBs could be explained by the highly proliferative state of aged HSCs, although not functionally competent, also noted in this study (10, 55). The transplanted restored cells may have changed the BM niche to provide HSCs with a functional microenvironment thereby changing age-linked secretome (56). Similar contribution of tissue niche in aging was reported for aged Paneth cells, resulting in intestinal stem cell dysfunction (57).

The aged donors are expected to have reduced clonal hematopoietic cell heterogeneity. The emerging clones in aged have an advantage due to mutations in identified genes (13). Our restoration system has not determined if there is enhanced clonal diversity, which in addition to emerging NK cells, could also ensure long-term health of the immune/hematopoietic system. Figure 7P summarizes the main findings, showing exosome with specific miR in co-cultures with young and old cells inducing partial reprograming of the aged cells. This is accompanied by increases in E2F/MYC and decreased p53. The reprogrammed hematopoietic cells showed changes in key aging hallmarks. More importantly, NK cells within the restored cells showed potential for immune therapy as indicated by dormant breast cancer clearance.

## Materials and Methods

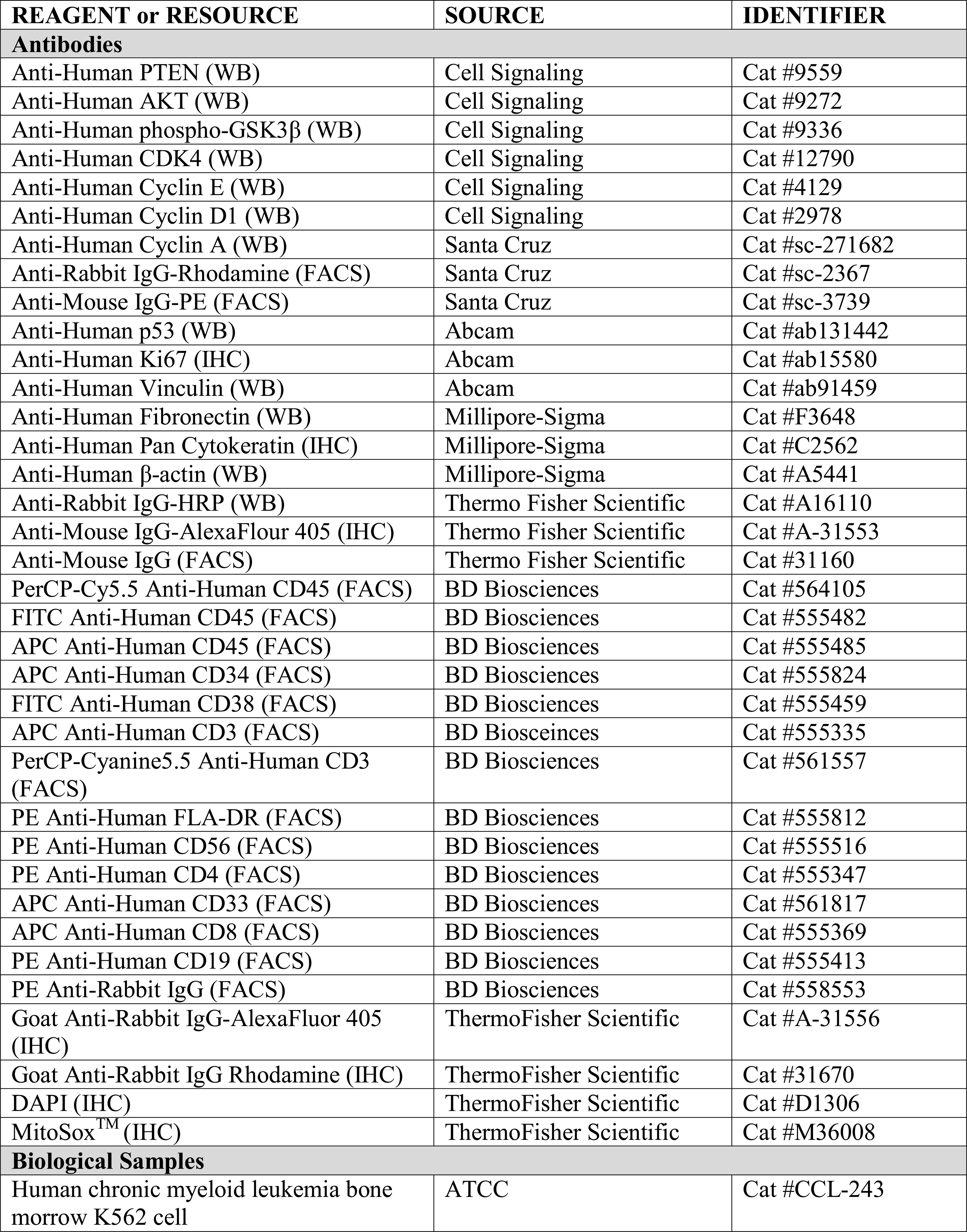
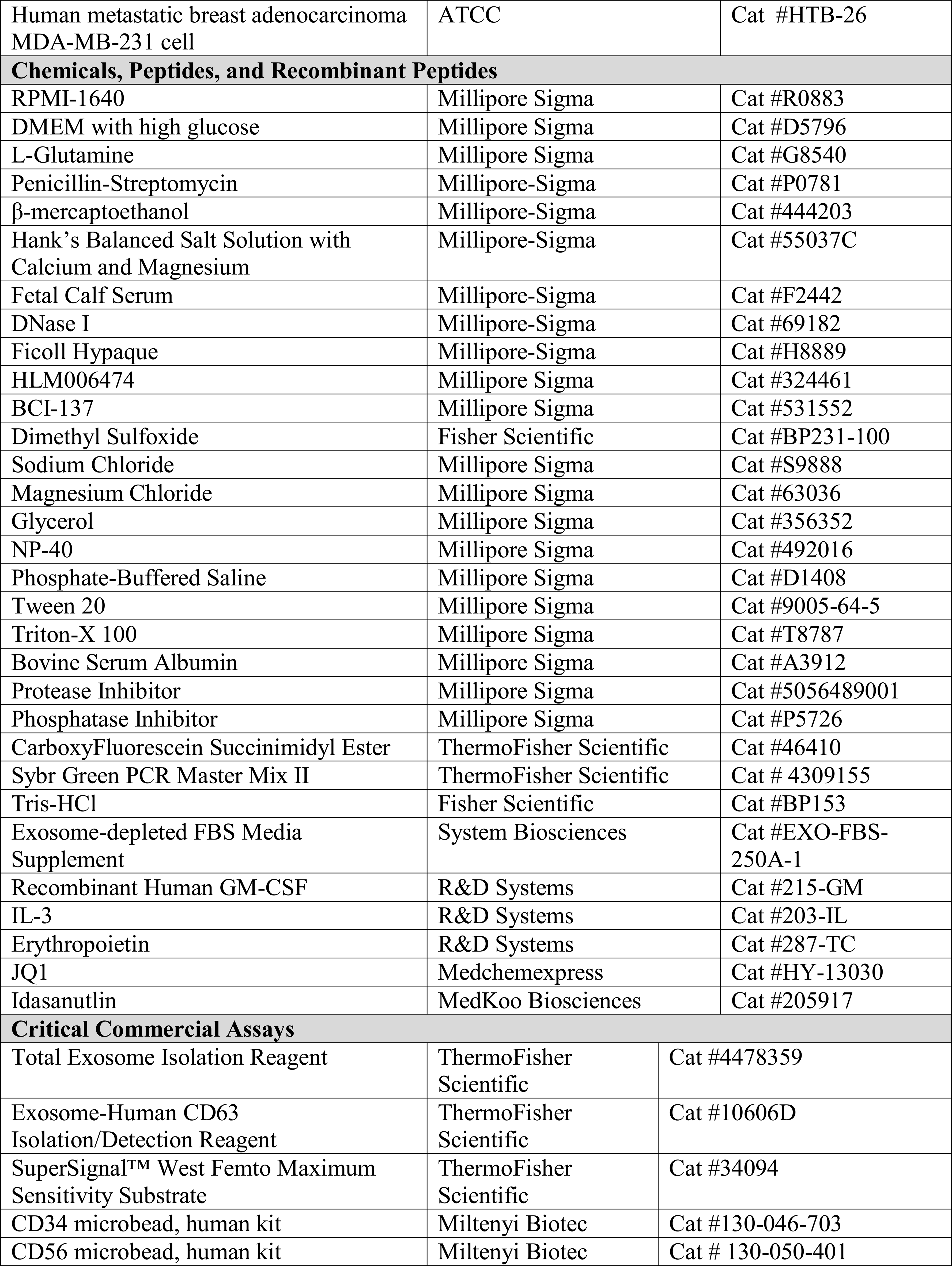
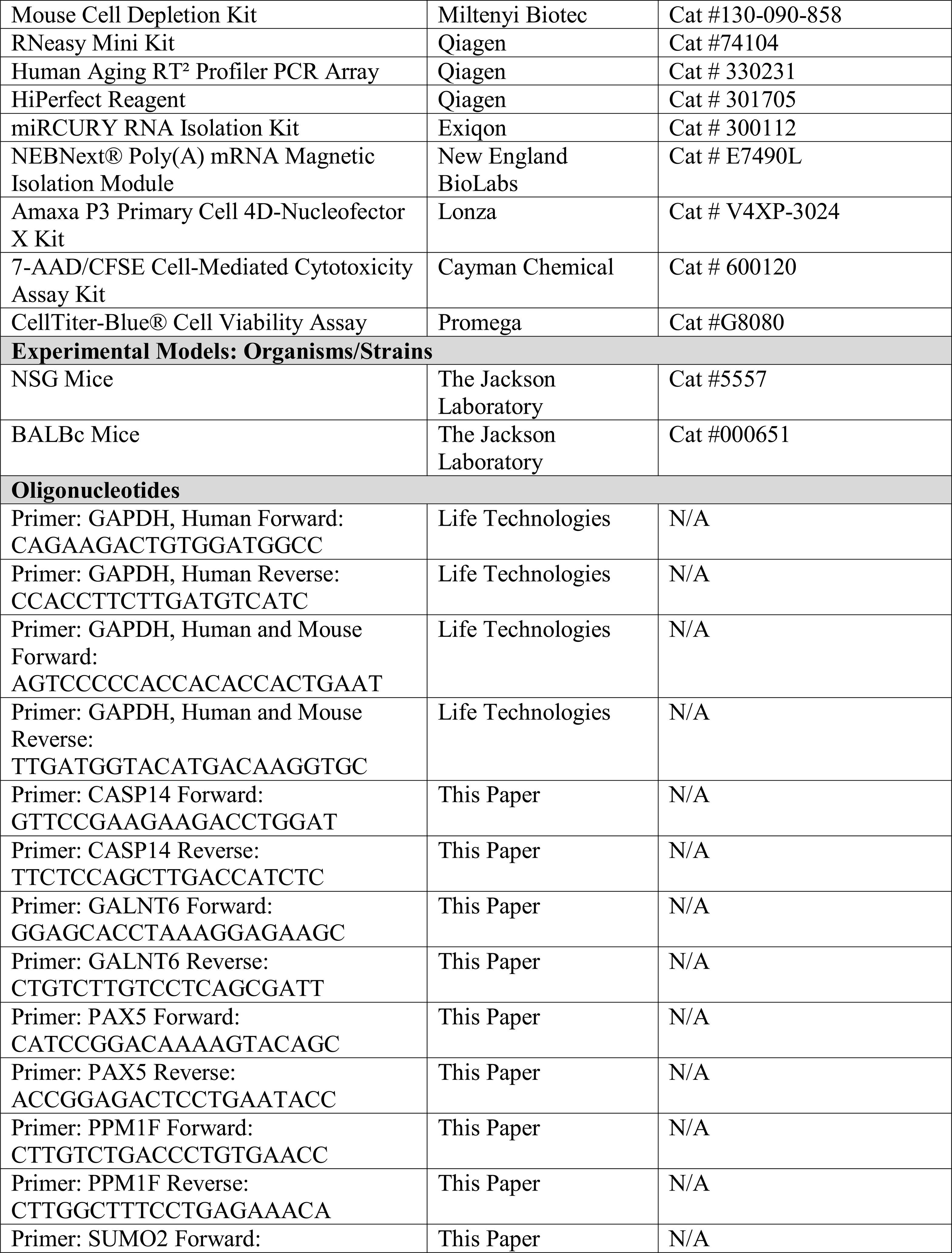
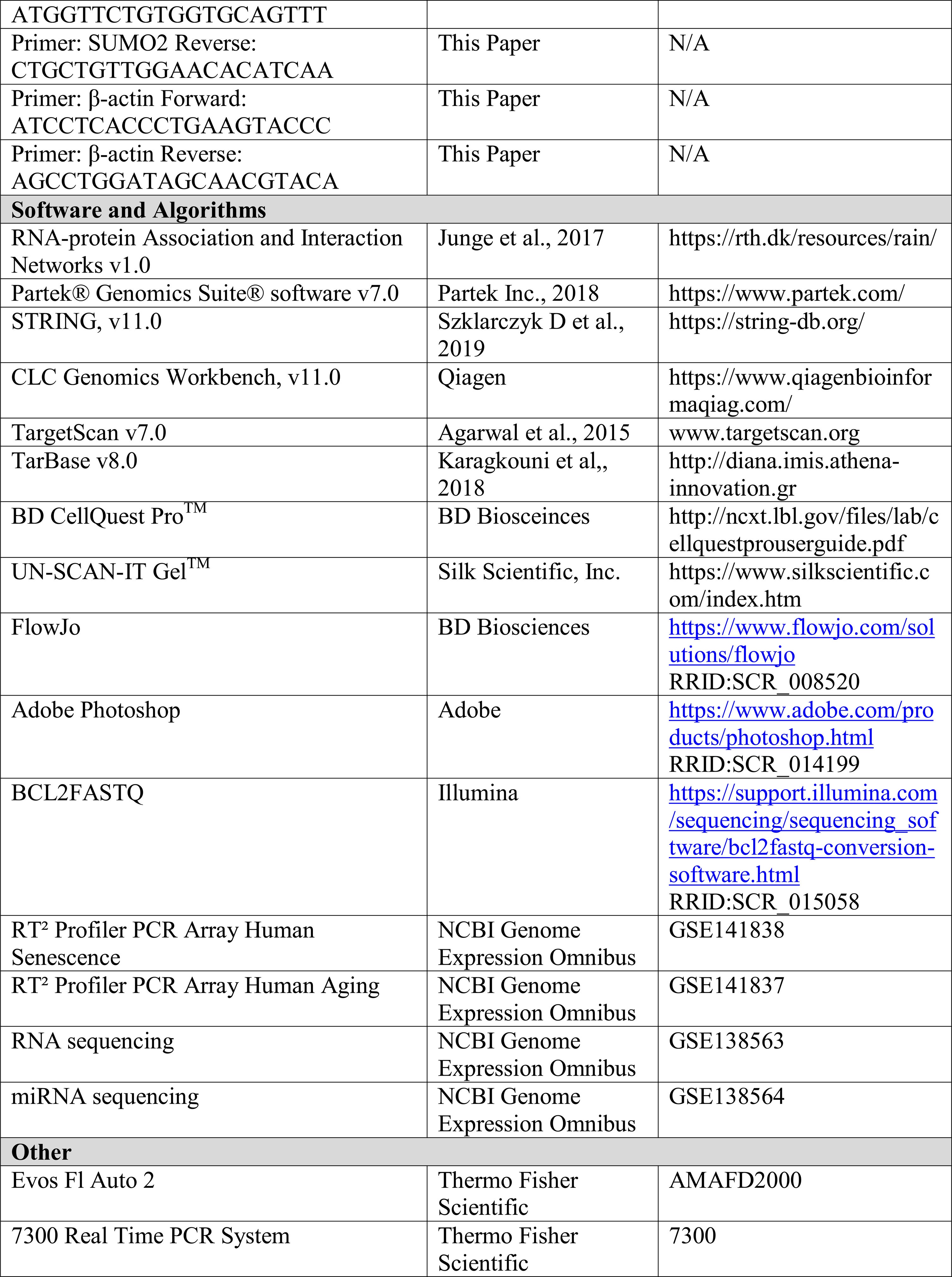
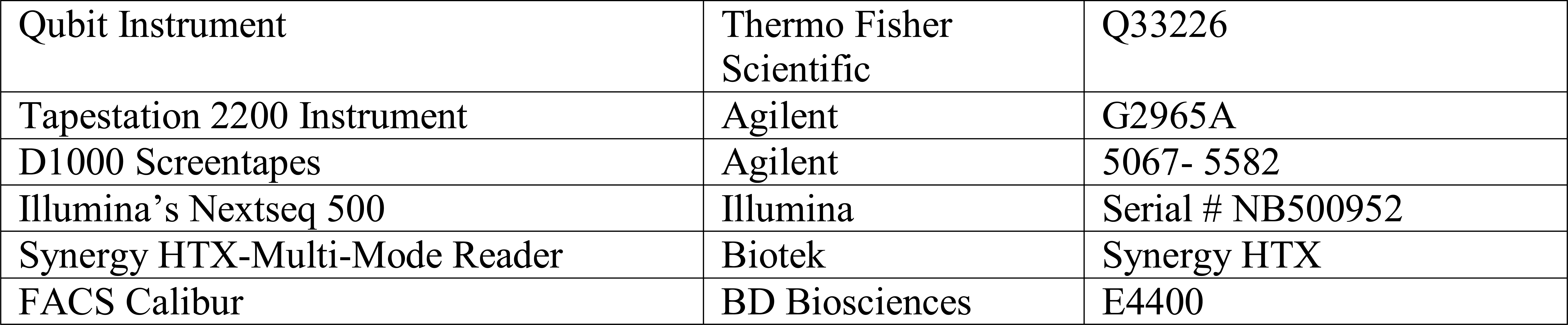

### Cell Lines/Validation

K562 and MDA-MB-231 were cultured as per American Type Culture Collection instruction. The cell lines were tested by Genetica DNA Laboratories (Burlington, NC) and were were verified as the original cells using ATCC STR database (www.atcc.org.org/STR_Database.aspx).

### Human Subjects

The use of de-identified human mobilized peripheral blood (MPB), peripheral blood (PB) and umbilical cord blood (UCB) was approved by Rutgers Institutional Review Board (IRB), Newark, NJ.

#### MPB

MPB was collected from aged (>60 yrs) and young (<30 yrs) donors (Table S1). Aliquots (10-20 mL) of MPB for A5-A8 (aged) and Y6-Y8 (young) were donated for research purposes through Progenitor Cell Therapy (Allendale, NJ). Donors A1-A4 (aged) and Y1-Y5 (young) were recruited, mobilized and subjected to leukapheresis by HemaCare Corporation (Van Nuys, CA). HemaCare was approved as a collection site by Rutgers IRB. The site is FDA-registered, AABB-accredited and operated under GMP-compliance. Study participants were given subcutaneous Neupogen® (G-CSF) at 5 μg/kg/day for 5 days. On day 6, MPB was collected using the Spectra Optia® Apheresis System to process 18 L of blood at a flow rate of 50 to 100 mL per min. Immediately after mobilization, the MPB were shipped to Rutgers.

#### Umbilical Cord Blood (UCB)

The demographics obtained for 10 UCB samples are shown in Table S1. The samples were provided within 24 h of collection by the Community Blood Services (Montvale, NJ), which is an AABB-accredited blood bank registered with the FDA. The mononuclear fraction was isolated by Ficoll-Hypaque density gradient and then cryopreserved for later use.

#### Peripheral Blood (PB)

PB was obtained from healthy donors (18-30 yrs) and the mononuclear fraction isolated by Ficoll Hypaque density gradient.

### Isolation of CD34^+^ Cells and Cell Cryopreservation

Total MPB, herein referenced as total nucleated cells (TNC), were cryopreserved by adding chilled cryopreservation media to the cell suspension at a 1:1 ratio while gently shaking the cells. The resuspended TNC was ∼ 5 x 10^7^ cells/mL. Cells were stored using a controlled rate freezer (Cryo Met Freezer, Thermo Fisher) at −1°C/min until the temperature reached −100°C. After this, the cells were transferred into liquid nitrogen.

CD34^+^ cells were selected using the CD34 microbead human kit (Miltenyi, Auburn, CA). The method followed manufacturer’s instructions. Briefly, TNC was pelleted at 4°C, 10 min, 500 *g*. The TNCs were resuspended in cold MACS buffer at a concentration of 10^8^ cells/300 µL buffer. Cell suspension was incubated in 100 µL of FcR Blocking Reagent and CD34 MicroBeads for 30 min at 4°C. After this, cells were washed with MACS buffer and the CD34^+^ cells were positively selected by magnetic separation with LS columns. The purified CD34^+^ cells were immediately cryopreserved as described above.

### Transwell Assay

Non-contact cultures were performed in 6- or 12-well transwell system, separated by a 0.4 μ membrane (BD Falcon, ThermoFisher). The inner wells contained 10^7^ young MPB or CB with the same number of old MPBs in the outer wells. The cells were cultured for one month in RPMI 1640 supplemented with 10% FCS, 2 mM glutamine and 0.5 µM β-mercaptoethanol (β-ME) (R10 media). Cultures with allogeneic cells were designated heterochronic and those with autologous cells, isochronic. During the culture period, 50% percent of media were replaced every four days with fresh R10 media.

### Cell Migration from Inner to Outer Wells

Transfer of cells from the inner to the outer wells of co-cultures were studied as follows: young MPBs were labeled with CarboxyFluorescein succinimidyl ester (CFSE) (Thermo Fisher) following manufacturer’s recommended protocol. Briefly, CFSE was diluted in 1x PBS to 5 μM (stock solution) and then diluted at 1/1000 (working solution). The latter was used to label cells by incubating for 20 min at room temperature. The labeled cells were added to the inner wells of the heterochronic cultures and at weekly intervals, the cells in the outer wells were analyzed for CFSE by flow cytometry.

### Flow Cytometry

Cells (10^6^) were labeled with the primary antibodies in 100 µl PBS. Each labeling used an isotype IgGs for background labeling (control) at the same concentration as the primary antibodies (see above for concentrations). After 30 mins at room temperature, the cells were washed with PBS and acquired on a FACSCalibur flow cytometer (BD Biosciences). Data were analyzed using BD CellQuest Pro™ software (BD Biosciences).

### Western Blot Analyses

Cell extracts were isolated with cell lysis buffer (50 mM Tris-HCl (pH 7.4), 100 mM NaCl, 2 mM MgCl_2_, 10% glycerol, and 1% NP-40) as described (Ghazaryan et al., 2014). Extracts (15 μg protein) were electrophoresed on SDS-PAGE and then transferred to polyvinylidene difluoride membranes (Perkin Elmer). The membranes were incubated overnight at 4^0^C on rocker with primary antibodies in 5% milk dissolved in 1x PBS tween. Next day, the membranes were incubated with species-specific HRP tagged secondary IgG in the same diluent. After 2 h, the membranes were developed by chemiluminescence using SuperSignal West Femto Maximum Sensitivity Substrate (Thermo Fisher Scientific).

### Clonogenic Assays

Clonogenic assays for hematopoietic progenitors, granulocytic-monocytic (CFU-GM) and early erythroid (BFU-E), were performed as described in methylcellulose matrices (Rameshwar et al., 2001). Cultures for CFU-GM contained 3 U/mL GM-CSF and BFU-E, 3 U/mL rhIL-3 and 2 U/mL Epo. Colonies were counted by a blinded observer. Each colony contained >15 cells.

### Modified Long-Term Culture-Initiating Cell (LTC-IC) Assay

Confluent stromal cells in the outer well of the transwell cultures were subjected to 1.5 Gy, delivered by a cesium source. After 16 h, the floating cells were washed and 10^7^ BM mononuclear cells added to the γ-irradiated stroma. At wks 6, 10 and 12, aliquots of mononuclear cells were assayed for CFU-GM in clonogenic assays (see above) (58).

### Mixed Lymphocyte Reaction (MLR)

One-way mixed lymphocyte reaction (MLR) was performed as described previously (59). Briefly, γ-irradiated (20 Gy) stimulator cells from isochronic or heterochronic cultures were added to 96 well-flat bottom plates (Corning, Corning, NY) at 10^6^/mL in 0.1 mL volume in triplicates. Autologous responder freshly thawed cells were added at the same concentration and volume. Control MLR used peripheral blood mononuclear cells (PBMCs) from two different donors (20-30 yrs). The MLR reactions were incubated at 37°C. At 72 h, the wells were pulsed with 1 μCi/well of [methyl-^3^H]TdR (70–90 Ci/mmol; NEN Radiochemicals, PerkinElmer, Akron, OH). After 16 h, the cells were harvested with a PhD cell harvester (Cambridge Technologies, Bedford, MA) onto glass-fiber filters, and [^3^H]TdR incorporation quantified in a scintillation counter (Beckman Coulter, Brea, CA). The results are expressed as the stimulation index (S.I.), which is the mean dpm of experimental cultures (responders + gamma-irradiated stimulators)/dpm of responder cells cultured in media alone.

### Cell Titer Blue Viability Assay

The cells in the outer wells of transwell cultures were assayed for viability with CellTiter-Blue Cell Viability Assay kit (Promega, Madison, WI). Viability analyses followed manufacturer’s instructions. Briefly, CellTiter-Blue reagent was added to aliquots of cells in 96-well plates. The plates were incubated for 4 h at 37°C and then read on a fluorescence microplate reader at 560 nm excitation/590 nm emission. Percent viability was calculated from a reference using untreated healthy cells, which were considered 100% viable, and cell-free wells containing reagent alone, which were considered 0% viable.

### Real time PCR for human cells in mouse femur

RNA extraction was performed according to manufacturer’s protocols with the RNeasy Mini Kit (Qiagen, Germantown, MD). Quality and concentration of RNA were determined with the Nanodrop ND-1000 spectrophotometer (ThermoFisher Scientific). The High-Capacity cDNA Reverse Transcription Kit (Life Technologies) was used to convert RNA to cDNA and 10 ng used in real-time PCR with Sybr Green PCR Master Mix II (Life Technologies) using primers from human and mouse GAPDH (see above).

### PCR array

Total RNA (2 μg) was extracted from MPB with the RNeasy Mini Kit (Qiagen) and then reverse-transcribed using the First Strand cDNA Synthesis Kit (Life Technologies). The cDNA (200 ng) was used for quantitative PCR with the Human Cellular Senescence RT^2^ Profiler™ PCR Array (Qiagen) containing 84 key genes involved in the initiation and progression of the biological process causing cells to lose the ability to divide. Prepared cDNA templates were added to the ready-to-use PCR master mix at equal volumes within each well of the same plate. The arrays were then run on the 7500 Real Time PCR System (Life Technologies) with the cycling profile (40 cycles) as follows: 94°C for 15 secs and 60°C for 45 secs. The data were analyzed with Qiagen PCR Array Data Analysis Software and normalized to five housekeeping genes provided within each array. Controls were also included for genomic DNA contamination, RNA quality, and general PCR performance. The normalized data are presented as fold difference, with a value of 1 representing no change. Differential and overlapping miRNAs are presented in Venn diagram. The overlapping areas include miRNAs with <1.5-fold difference among groups.

### Exosome Isolation and Nanoparticle Tracking Analysis

Exosomes were isolated from cell culture media as described (Bliss et al., 2016). In addition, exosomes were also isolated with the Total Exosome Isolation Kit (Life Technologies), using a modified version of the manufacturer’s protocol. Specifically, isochronic and heterochronic cultures were established with Exosome-depleted FBS Media Supplement (System Biosciences, Palo Alto, CA). Culture media collected on the 4^th^ and 7^th^ days were pelleted and supernatant transferred to another tube for further clarification at 2000 *g* for 30 min to remove residual cells and debris. The remaining supernatant was transferred to a fresh tube and 0.5 volumes of Total Exosome Isolation reagent was added for overnight incubation at 4°C. The following day, samples were centrifuged at 10,000 *g* for 1 h at 4°C to pellet the exosomes for subsequent nanoparticle tracking analysis (NTA) or long-term storage at −80°C. The absolute size distribution of exosomes was performed with the NanoSight LM10 and the NTA3.1 software (Malvern Panalytical, Malvern, UK). The particles were automatically tracked and sized, based on Brownian motion and diffusion coefficient. NTA analyses resuspended the exosomes in 0.5 mL PBS with the following parameters: Temperature = 25.6 +/− 0.5°C; Viscosity = (Water) 0.99 +/− 0.01 cP; Measurement time = 30 sec; Syringe pump speed = 30. The detection threshold was similar in all samples. We record the data from each sample thrice.

### MitoSox Assay

MPB cells (10^6^) were labeled with anti-CD34-APC and -CD45-PerCp-Cy5.5, as described below for flow cytometry. The cells were washed and incubated with 5 μM MitoSox™ Red for 10 min at 37°C in the dark and then washed again with warm HBSS. The data was acquired and analyzed with the FACSCalibur.

### Senescence Associated Secretory Phenotype (SASP) array

#### Media

Senescence associated secretory factors (SASFs) in media from heterochronic cultures at day 1 and wk 4 were analyzed with Custom C-Series Human Antibody Arrays with 68 different factors linked to cellular senescence (60). The antibodies were obtained from Ray Biotech (Norcross, GA). Briefly, cell culture media were pelleted at low speed centrifugation (300 *g*) to remove cell debris and stored in siliconized microfuge tubes at −80°C until assayed. Incubation and detection of factors within the conditioned media followed the manufacturer’s suggested protocol. Background levels were calculated by incubating the arrays with complete media alone and then subtracting the obtained values from each conditioned media experiment. Densitometry was performed using the UN-SCAN-IT densitometry software (Silk Scientific; Orem, UT).

#### Plasma

Detection of SASFs in plasma of huNSG mice was performed as described above for media. Briefly, blood from huNSG mice was pelleted for 10 min at 300 *g* and the plasma supernatant collected in siliconized microfuge tubes for SASF determination. Background was subtracted from the density of parallel analyses with plasma from non-humanized NSG mice.

### Senescence and Aging Gene Arrays

Cells were flushed from the femurs of huNSG using a 26-gauge needle. Murine cells were eliminated with a Mouse Cell Depletion Kit. Total RNA (2 μg) was extracted from purified cells using the RNeasy Mini Kit and reverse-transcribed with the RT^2^ First Strand Kit. 20 ng of cDNA was used for qPCR with the Human Cellular Senescence and Human Aging RT^2^ Profiler™ PCR Arrays (Qiagen). Arrays were run on the 7300 Real Time PCR System (Life Technologies) with the cycling profile (40 cycles): 94°C for 15 secs and 60°C for 45 secs. Gene expression analysis was performed using Qiagen PCR Array Data Analysis Software. The data were normalized to five housekeeping genes provided within each array (http://pcrdataanalysis.sabiosciences.com/pcr/arrayanalysis.php). The data was submitted to the NCBI Genome Expression Omnibus (GEO; https://www.ncbi.nlm.nih.gov/geo/) under SuperSeries accession number GSE138565. Hierarchical clustering and heat map generation were performed with Heatmapper software (http://www.heatmapper.ca/), as described (61).

### MiRNA microarray and qPCR

Total RNA (500 ng) was isolated from exosomes using the miRCURY RNA Isolation Kit (Exiqon). RNA was reverse-transcribed with the miScript II RT Kit (Qiagen) and 20 ng of cDNA used for qPCR with the human miFinder miRNA Array (Qiagen). The PCR was done with the following cycling conditions: 94°C for 15 mins, 40 cycles at 94°C for 10 secs, 55°C for 30 secs, 70°C for 30 secs, followed by melt curve analysis. The data were analyzed with the online miScript miRNA PCR Array data analysis tool (Qiagen).

Validation of miRNAs identified in the microarray and NGS was done by individual qPCR experiments using miScript primer assays and similar cycling and analysis schemes. Total RNA (2 µg) was also isolated from cells, as described above, for profiling of downstream miRNA targets by qPCR using the primer pairs listed above with the following cycling conditions: 95°C for 15 mins, 40 cycles at 94°C for 15 secs, 51°C for 30 secs, 72°C for 30 secs, followed by melt curve analysis. Analyses were performed with Qiagen PCR Array Data Analysis Software, as described above. Array and individual qPCR studies were normalized to RNU6, SNORD68 and SNORD95 and presented as fold change. The 30 genes with marked changes following restoration were selected for links to other genes using RNA-Protein Association and Interaction Networks (RAIN) (62).

### RNA Sequencing (Seq)

RNA Seq was performed at the Genomics Center at Rutgers New Jersey Medical School (Newark, NJ). Total RNA was submitted to the center where poly A RNA was purified using NEBNext® Poly(A) mRNA Magnetic Isolation Module (New England BioLabs, Ipswich, MA). Next generation sequencing (NGS) libraries were prepared using the NEB Ultra II Library Preparation Kit and NEBNext® Multiplex Oligos for Illumina (Dual Index Primers Set 1) (New England BioLabs). The generation of the libraries followed manufacturer’s protocol. The libraries were subjected to quality control using Qubit instrument and high sensitivity Kit from Thermo Fisher as well as Tapestation 2200 instrument and D1000 ScreenTapes from Agilent (Santa Clara, CA). The libraries were diluted to 2 nM and then denatured as per Illumina’s protocol and run on Illumina’s NextSeq instrument (San Diego, CA) using 1X75 cycle high throughput kit. The BCL files that were generated from the sequencing were demultiplexed and converted to FastQ files using BCL2FASTQ software from Illumina. All raw and processed sequencing data have been submitted to the NCBI Genome Expression Omnibus (GEO; https://www.ncbi.nlm.nih.gov/geo/) under accession number GSE138563, SuperSeries accession number GSE138565.

### MiRNA sequencing (miR Seq)

Total RNA from exosomes and cells was isolated using the miRCURY RNA Isolation Kit with small and large RNAs fractionated with the RNeasy MinElute Cleanup Kit, both according to manufacturer’s recommended specifications (Qiagen). Half of the small RNA fraction (200 ng) was used in library preparation with the NEBNext Multiplex Small RNA Sample Prep Set for Illumina – Set 1 (New England Biotechnology), according to the following protocol: (1) ligation of the 3′ SR Adaptor, (2) hybridization of the reverse transcription primer, (3) ligation of the 5′ SR Adaptor, (4) reverse transcription for first strand cDNA synthesis and (5) PCR enrichment. After PCR, samples were cleaned and size selection performed. Briefly, 2 μl of sample was subjected to TapeStation (Agilent) analysis to ascertain band sizes. Samples were run on 8% acrylamide gel at 100V for 1 hr, with correct size bands excised for gel purification. Small RNA libraries were diluted to 2 nM and run on a miSeq System (Illumina) for NGS using the V2 kit (Illumina). All raw and processed sequencing data have been submitted to the NCBI Genome Expression Omnibus (GEO; https://www.ncbi.nlm.nih.gov/geo/) under accession number GSE138564, SuperSeries accession number GSE138565.

### Data Analyses

#### RNA Network

Specific genes from the senescence array data were analyzed for RNA interaction using the RNA-protein Association and Interaction Networks (v1.0) (RAIN) (62). The upregulated genes following restoration were input into the program. The database projected the output in the form of networks and co-expression graphs.

#### RNA Seq

Partek^®^ Flow^®^ software was used to align the reads to Homo Sapiens assembly hg38. Transcript abundance of the aligned reads were performed with Partek’s (version 7) optimization of the expectation-maximization algorithm (Partek Inc., St. Louis, MO). The data trends and outliers were detected by Principle Component Analysis (PCA). The RNA-Seq data was normalized by using log_2_ counts per million (CPM) with an offset of 1. Batch effect was removed between two sequencing rounds of the same biological sample. Differential analyses were performed using Partek™ GSA with an FDR q-value< .05 and a fold change of 1.3 in aged vs. restored, 1.45 in Restored vs. Young, and 1.65 in Aged vs Young. The fold change cutoff was based on point of maximum curvature in the ranked fold changes in each comparison. The analyses resulted in 2,140 genes. The differences in gene expression were visualized by hierarchal clustering using Morpheus. Clusters were then identified by visual analysis as well as dendrogram branches. Top gene ontology (GO) terms for each cluster were found using STRING (Version 11) and the significant pathways and functions were displayed next to the hierarchal clustering (63). Additionally, pathway and functional analyses were done by Gene Set Enrichment Analysis (GSEA). False discovery rate of ≤0.05 was used as a cutoff to select significantly up or downregulated pathways.

#### Ingenuity Pathway Analysis (IPA) – Hematological Effects Analysis

Pathways of genes linked to hematological functions were selected using Ingenuity Pathway Analyses (IPA, QIAGEN Inc.). These pathways were ranked by Significance score, defined as −Log_10_B-H *p* value times activation z-score. We retained the pathways associated with hematological functions that passed a significance score threshold, >2.

#### MiR Seq

Data analyses were performed using CLC Genomics Workbench (Qiagen) according the following data workflow: (a) Fastq files were imported into the analysis suite, (b) sequences were trimmed to remove poor quality and short reads, (c) trimmed reads were run through the Small RNA Analysis pipeline, (d) extraction and counting, (e) annotation and count merging to identify expression level of each mapped miRNA. Mapped reads from individual samples were then compared to determine fold change for each miRNA.

#### MiRNA Target Prediction and Network Analyses

All miRNAs exhibiting greater than 100 mappable reads were analyzed. Expression data from NGS was analyzed in silico by IPA to predict miRNA targets and downstream signaling networks. Differentially expressed exosomal and intracellular miRNA (1.4-fold cutoff) between young and aged isochronic, and aged isochronic and heterochronic samples were uploaded to the IPA suite for Core Network Analysis. Predicted networks from the Core Analyses were then simultaneously linked using Comparison Analyses to identify the exosome-cell interactome during heterochronic restoration. Potential mRNA targets of candidate miRNAs were determined using the miRNA Target Filter. The source of the miRNA-mRNA relationship and the confidence of the relationship predictions were from TargetScan (www.targetscan.org) and the experimentally observed relationships were from TarBase (http://diana.imis.athena-innovation.gr). mRNA target selection was based on target rank score, where the highest ranked targets were common to the most candidate miRNA (score = 6) and the lowest ranked targets to the least candidate miRNA (score = 1). Potential interaction with the exosome – cell interactome was evaluated by creating a mock mRNA target expression profile (10-fold downregulation) to generate a Core Analysis network that could be likened using the Comparison Analysis tool. Candidates whose predicted networks converged with the interactome were selected for additional evaluation.

#### Targets of Validated miRNAs

Analyses of 6 miRNAs for predicted targets were performed with TargetScan human database. A total of 6101 potential targets were evaluated, with a number of common targets within the group of 6 miRNA displayed within the descending concentric circles. 25 targets were identified that met the following conditions: (1) ≥ 4 common hits among the miRNA group, including miR-619 OR miR-1303 or, (2) ≤ 3 common hits among the group, including miR-619 and miR-1303. Predicted expression of these targets was analyzed by IPA and the resulting network predictions compared to the young exosomal and aged heterochronic intracellular miRNA interactome. We selected the tabulated targets by paring down based on their expression in relevant tissues. Specifically, if they encode verified protein and, their expression is not limited only in neural tissue.

qPCR miRNA Array Analyses: Total RNA (500 ng) was isolated from exosomes using the miRCURY RNA Isolation Kit and reverse-transcribed with the miScript II RT Kit. 20 ng of cDNA was used for qPCR with the human miFinder miRNA Array, with cycling conditions of 94°C for 15 min, 40 cycles at 94°C for 10 seconds, 55°C for 30 secs, 70°C for 30 secs, followed by melt curve analysis. The data were analyzed with the online miScript miRNA PCR Array data analysis tool (SABiosciences). The miRNAs identified by microarray and NGS were validated by real time PCR with miScript primer assays using similar cycling and analysis schemes. Total RNA (2 µg) was also isolated from cells, as described above, for profiling of downstream miRNA targets by real time PCR using primers listed above and the following cycling conditions: 95°C for 15 mins, 40 cycles at 94°C for 15 secs, 51°C for 30 secs, 72°C for 30 secs, followed by melt curve analysis. Analyses were performed with Qiagen PCR Array Data Analysis Software, as described above.

### Nucleofection of miRNA Mimics, miRNA Inhibitors and siRNA

Aged TNCs (10^7^/sample) were nucleofected with miRNA mimics (Qiagen), miRNA inhibitors (Qiagen), negative control siRNA (Qiagen), negative control miRNA inhibitor (Qiagen) or downstream target candidate siRNAs (Origene) using the Amaxa P3 Primary Cell 4D-Nucleofector X Kit (Lonza) on a 4D Nucleofector device (Lonza), according to manufacturer’s specific protocol. Briefly, CD34^+^ cells were nucleofected with 60 nM of total miRNA mimics, miRNA inhibitors or siRNA using the “human CD34^+^ cell” program.

### Sorting of Breast Cancer Stem Cells (CSCs)

MDA-MB-231 breast cancer cells were stably transfected with pEGFP1-Oct3/4, as previously described (50). The CSCs were selected as the top 5% cells with the highest GFP intensity (Oct4^hi^) and then sorted with the FACSAria II cell sorter (BD Biosciences).

### *In Vivo* Studies

The use of mice was approved by Rutgers Institutional Animal Care and Use Committee (Newark Campus, NJ). Mice were housed in an AAALAC-accredited facility.

#### Humanization (huNSG)

Female NSG mice (NOD.Cg-*Prkdcscid Il2rgtm1Wjl*/SzJNOD-*scid* IL2Rgamma^null^, 5 wks) were obtained from Jackson Laboratories (Bar Harbor, ME). These mice lack the major immune cells such as T-cells, B-cells, macrophages and NK cells. After 1 wk acclimatization, NSG mice were subjected to 150 cGy whole body γ-irradiation using a Mark-I cesium irradiator unit. At 5 h post-irradiation, the mice were injected intravenously with 5 x 10^5^ human CD34^+^ cells isolated from aged or young MPBs. At 9 and 13 wks post-transplantation, we assessed the mice for chimerism by flow cytometry for human CD45. Blood cells were co-labeled with anti-human CD45-APC and anti-mouse CD45-FITC.

#### *In Vivo* Restoration

huNSG with aged CD34+ cells were transplanted by intravenous injection of 5x10^5^ autologous aged cells from 7-day CD3-depleted isochronic (non-restored) or heterochronic (restored) cultures. Control mice were injected with sterile PBS. % huCD45+ cells were calculated from total nucleated cells; % huCD34+ cells were calculated from total huCD45+ cells.

#### Serial Transplant

Mice were injected with 10^5^ huCD34+ cells taken at the end point of the first set of transplants from the three groups.

#### MiRNA-based *In Vivo* Restoration

The miRNA-based studies were performed as above, except for a second injection with 60 nM miR-619, miR-combo (miR-619, -1303, 4497) or control miR. The cells were transfected with HiPerFect reagent and then cultured for 7 days. CD3+-depleted cells were transplanted as above. At the end-point of 14-15 wks post-transplant, blood, femurs and spleens were harvested for biochemical, phenotypic and functional analyses. Major organs were also harvested for histological assessment. Tissue embedding, processing and staining were performed by the Digital Imaging and Histology Core of Rutgers-New Jersey Medical School Cancer Center (Newark, NJ). Histologic findings were confirmed on H&E slides by a board-certified veterinary pathologist.

#### NK function in mice with breast cancer stem cell (CSC)

Nude female BALB/c mice (6 wks) were injected with CSCs for established dormancy in femurs as described (50). The CSCs were isolated as outlined above. Mice with dormant cancer cells were injected with 10^6^ restored or unrestored aged MPBs. After one month, mice were studied for human chimerism and GFP cells were analyzed by real time PCR for GFP. Groups of 10 mice were followed for one year for survival.

The role for NK cell in targeting dormant breast cancer cells in mice was studied with NSG due to the lack of endogenous NK cells. The method followed procedure used for nude BALB/c mice.

### Immunohistochemistry

Immunohistochemistry for human pan cytokeratin and Ki67 in murine femurs was performed as described (46). Briefly antigen was retrieved from deparaffinized sections of mouse femurs by heating at 56^0^C overnight. After this the slides were dewaxed with xylene and ethanol then rehydrated, fixed and permeabilized with 0.1% Triton X. Slides were then washed 3x in 1x PBS and then incubated overnight at 37^0^C in a humidified chamber with primary antibodies at 1/500 final dilution for anti-cytokeratin and anti-Ki67. The slides were washed and then incubated (2 hrs at room temp) in a humidified chamber with fluorescence-tagged secondary antibodies at 1/1000 final dilution: AlexFluor 405 for cytokeratin and rhodamine for Ki67. The slides were washed then covered with 1x PBS and then immediately analyzed on the EVOS FL Auto 2 Imaging System (ThermoFisher Scientific). Tissues from scraped femurs were placed on slides and then similarly imaged.

### Co-culture between natural killer (NK−/+) and CSCs

NK cell depletion in the restored and unrestored MPBs was conducted with human CD56 microbeads from Millitenyi (Auburn, CA). The method followed manufacturer’s instruction. Co-cultures were performed with 10^5^ breast CSCs and 10^7^ restored MPB, with or without NK cells. After 24 h, the number of CSCs were determined by the number of GFP+ breast cancer cells using fluorescence microscopy (Evos FL2 auto).

### NK Cell Assay

#### ^51^Cr release

NK cell function was determined by the [^51^Cr] radionuclide assay for cytotoxicity, which was performed as described previously (64). Briefly, the K562 cell line was used as targets (T), and MPBs as effector (E) cells. K562 cells (5×10^6^/ml) were labeled with 200 μCi of ^51^Cr (PerkinElmer, Wellesley, MA). After labeling, the K562 cells were washed three times and then resuspended at 10^5^/mL in RPMI 1640 + 10% FCS. The effector MPBs were resuspended at 10^7^/mL RPMI 1640 + 10% FCS. Equal volumes (100 μl) of effector and target were added to round bottom 96-well Corning plates (Millipore Sigma) in quadruplicates at E:T ratios of 100:1, 50:1, 25:1 and 12.5:1. Spontaneous [^51^Cr] release was determined by incubating targets alone, and total release was determined in parallel incubations with 1% Triton X-100. After 4 h of incubation, cell-free supernatants were collected, and the amount of [^51^Cr] release was determined in a γ-counter. The percentage lysis were calculated as follows: [experimental point (dpm) − spontaneous release (dpm)]/[total release (dpm) − spontaneous release (dpm)] × 100. Values of spontaneous release were <1% of total release. The average technical replicates were plotted on a graph of E:T vs % Lysis. The lytic units (LU) was determined as described (Bryant et al., 1992). One LU was assigned as the number of effector cells that lysed 20% of the target cells. Total LU is the number in 10^7^ effectors. Each replicate was used to calculate the total LU then plotted on the same experimental point as mean±SD.

#### Carboxyfluorescein succinimidyl ester (CFSE/7-AAD)

Cytotoxicity assay was performed with CFSE/7-AAD Cell Cytotoxicity kit (Cayman Chemical, Ann Arbor, MI), following manufacturer’s instructions. K562 cells (10^7^) were labeled with CFSE dye for 15 mins and then washed twice. Cells were diluted to 10^5^/ml and incubated for 30 mins at 37 °C. Day 7 cultures of restored or unrestored aged MPBs were used as effector cells. The effector and target cells were added into 12-well plates at the following E:T ratios: 0:1, 6.25:1, 12.5:1 and 25:1. The plates were incubated for 4 h at 37 °C. Cells were harvested and stained with 7-AAD. Events (50,000) were acquired on FACSCalibur flow cytometer. Data were analyzed using Cellquest pro software (BD Biosciences). Target cells stained only with 7-AAD were used to calculate spontaneous lysis while effector cells stained only with 7-AAD were used to detect dead effector cells. The % lysis was calculated as [Cells positive for both CFSE and 7­AAD/Total CFSE labelled cells] * 100 − spontaneous lysis.

### T-Cell Activation Assay

T-cell activation was determined using the T-Cell Activation/Expansion Kit. Briefly, anti-biotin MACSiBead Particles were loaded with CD2, CD3, and CD28 antibodies. Cells from 7-day isochronic or heterochronic cultures, or MNCs isolated from huNSG mouse blood by Ficoll-Hypaque density gradient, were incubated with loaded anti-biotin MACSiBead Particles at a 1:2 bead to cell ratio for 72 h to activate T-cells. Addition of unloaded MACSiBead Particles served as negative control. After 72 h, cells were fluorescently labeled using CD45-FITC, CD4-PE, CD25-PerCP-Cy5.5 and CD8-APC to determine T-cell activation status by flow cytometry.

### Statistical Analyses

Statistical analyses were performed with ANOVA and Tukey-Kramer multiple comparisons test. For array and NGS expression analyses, average linkage was used for clustering and Pearson correlation analysis used for distance measurement to generate heatmaps and hierarchically cluster genes. *p*<0.05 was considered significant.

## Acknowledgments

This work was supported by a grant from the Bosarge Family Office.

Thanks to Professor Rakesh Kumar for critical input on the manuscript.

## Author contributions

The work was conceptualized by SJG and PR; All authors were involved in conducting the experiments; SJG submitted the original draft; ME, OAS, LSS, RJD and SA analyzed the sequencing data; All authors edited the manuscript.

