## Supplemental Table 1 and Figures 1-6 for "Exosome-mediated hematopoietic rejuvenation in a humanized mouse model indicate potential for cancer immunotherapy"

**SUPPLEMENTAL INFORMATION**

**Title:** Recapitulation of hematopoietic rejuvenation by young cells in humanized mice indicate potential for cancer immunotherapy

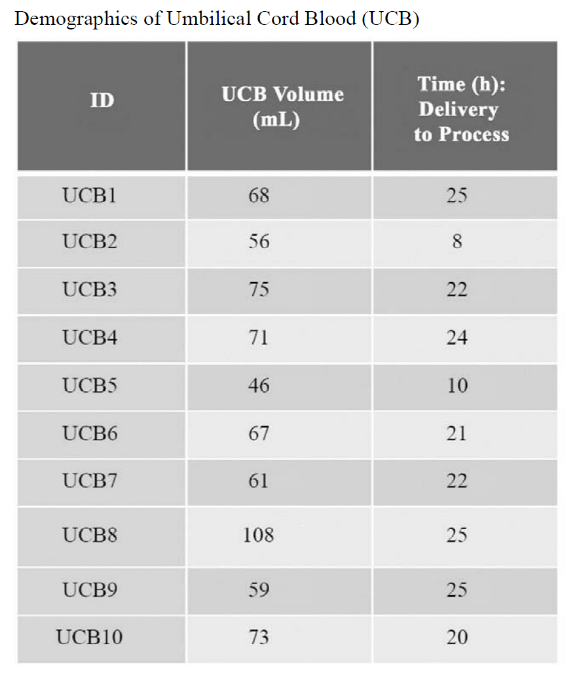

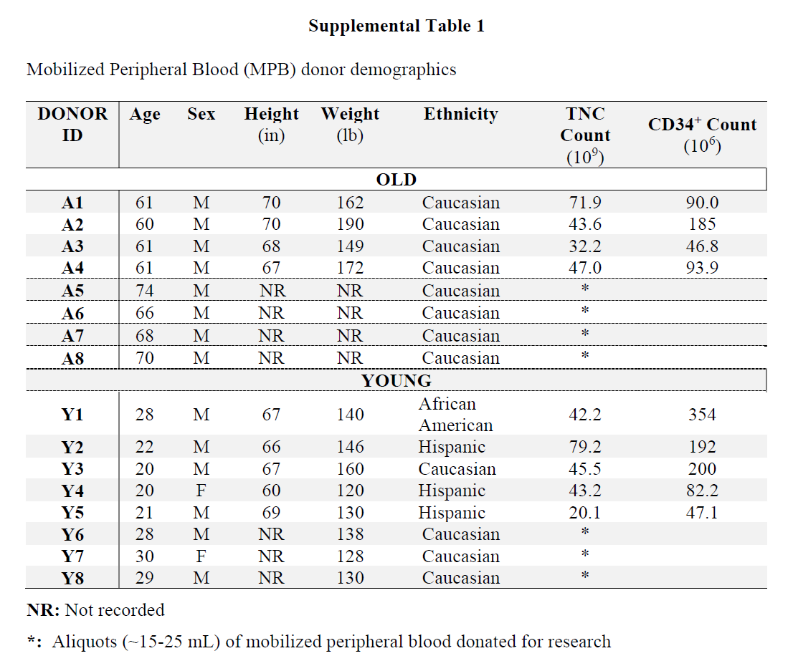
**Authors:** Steven J. Greco, Seda Ayer, Khadidiatou Guiro, Garima Sinha, Robert J. Donnelly^3^, Markos El-Far, Sri Harika Pamarthi, Oleta A. Sandiford, Marina Gergues, Lauren S. Sherman, Michael J. Schonning, Jean-Pierre Etchegaray, Nicholas M. Ponzio, Narayanan Ramaswamy, Pranela Rameshwar

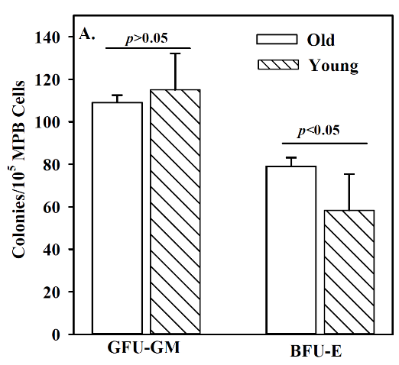

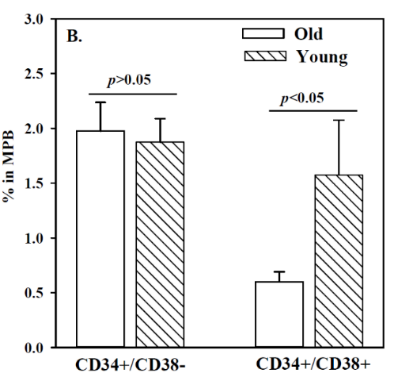

**C.**

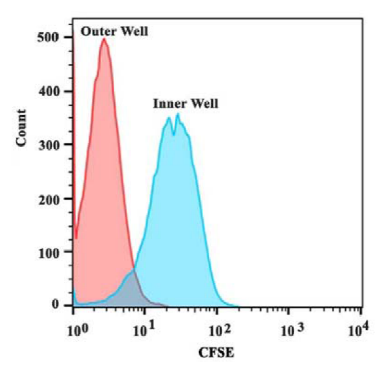

**D.**

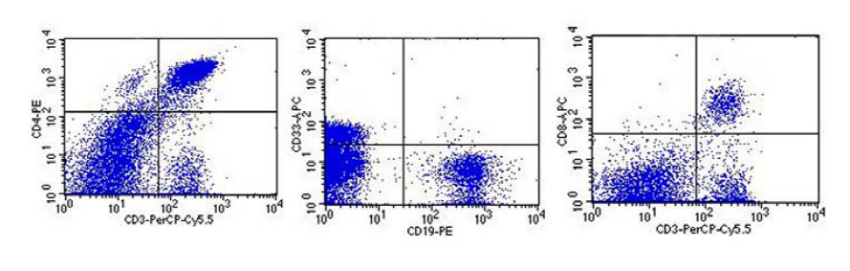

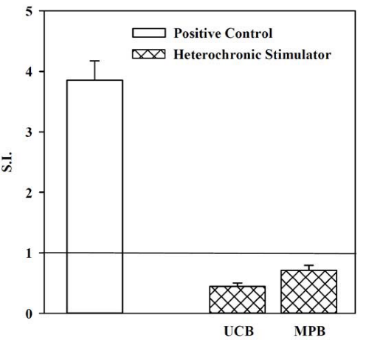

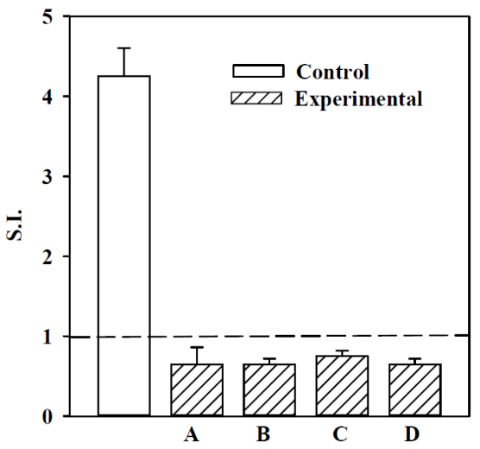

**D.**

**G.**

**E.**

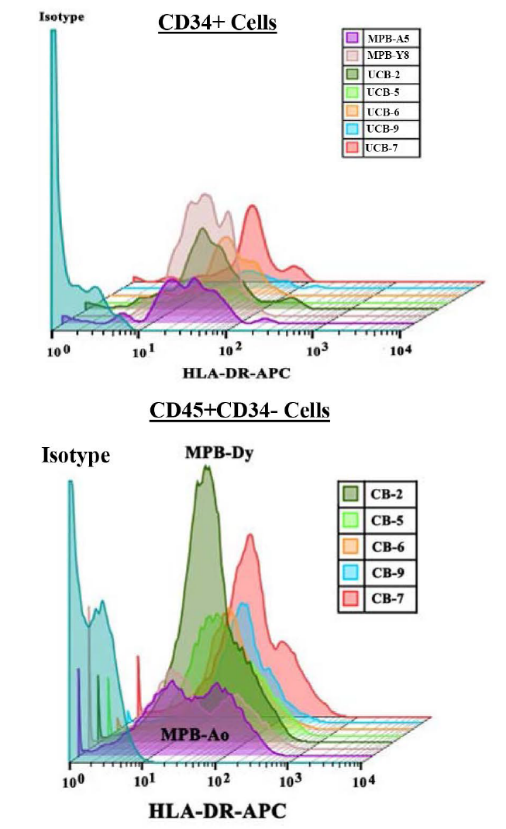

**F.**

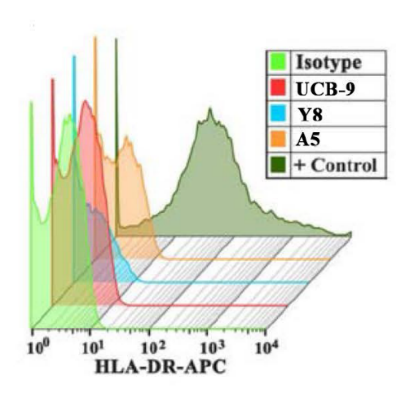

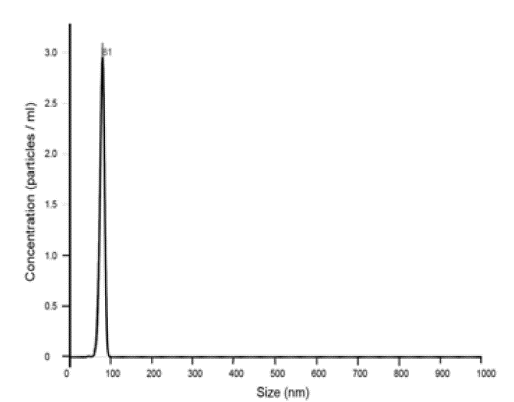

**I.**

**H.**

**Figure S1. (A)** Clonogenic assays for CFU-GM and BFU-E were performed with MPB from young and old donors (n=8 for each group, each tested in triplicate). CFU-GM/10^5^ cells are presented as the mean±SD. **(B)** Flow cytometry with cells from donors was performed by gating the CD34+ cells and then examining for CD38. The results are shown as the mean MFI±SD for four old and young donors. **(C.)** Transwell cultures were performed with CFSE-labeled young MPBs in the inner wells and unlabeled old MPBs in the outer wells. The inner wells were examined for CFSE weekly up to 4 wks. Shown is the histogram at the 4-wk time point. **(D.)** Shown are representative images of flow cytometry for T- (CD3 and CD4), B- (CD19) and myeloid (CD33) cells in old MPBs. **(E)** MLR with old restored MPB as stimulator and naïve autologous old MPB as responder: Heterochronic cultures were established with MPB from four old donors (arbitrarily labeled A-D) and two different young MPBs. After 4 wks, the restored old MPBs were γ-irradiated and then used in a one way MLR as stimulator cells. Freshly thawed autologous MPBs were used as responders. Positive control used unrestored allogeneic young or old MPBs. The results are presented as mean S.I.±SD for each donor, each tested with two young MPBs in duplicate. The positive controls, which represented 4 experimental points, two with allogeneic old MPBs and 2 with allogeneic old MPBs, showed similar outcomes and were plotted on the same bar. Each MLR test was performed in quadruplicate. **(F)** Flow cytometry for MHC-II using CD34+ cells (top panel) and CD34-/CD45+ (bottom panel) from 5 randomly selected UCB. Comparison analyses used MPBs from young and old donors. The cells were labeled with anti-HLA-DR-APC or isotype. The results show one representative young and old MPB. The results indicated higher density of MHC-II on CD34+ and CD34- UCB as compared to MPB from an old and a young donor. **(G.)** MLR reactions were repeated as above, except for heterochronic cultures containing UCB in the inner wells. The results are shown as the mean S.I.±SD from five heterochronic cultures in which four old MPBs were restored with five different UCB. Each heterochronic culture was done in duplicate. The MLR reactions were performed in quadruplicates. **(H.)** Representative histogram of exosomes analyzed isolated from the media of heterochronic cultures. The analysis was done on nanotracking analyzer (Nanosight, Malvern, Westborough, MA). **(I.)** Exosomes were captured onto CD63-coupled beads and then analyzed by flow cytometry for MHC-II. The beads were labeled with anti-HLA-DR-APC. Representative results are shown for exosomes of heterochronic cultures performed with UCB. Parallel analyses were performed with media from isochronic cultures containing young and old MPBs. Positive control (+ control) used anti-CD3 activated mononuclear cells.

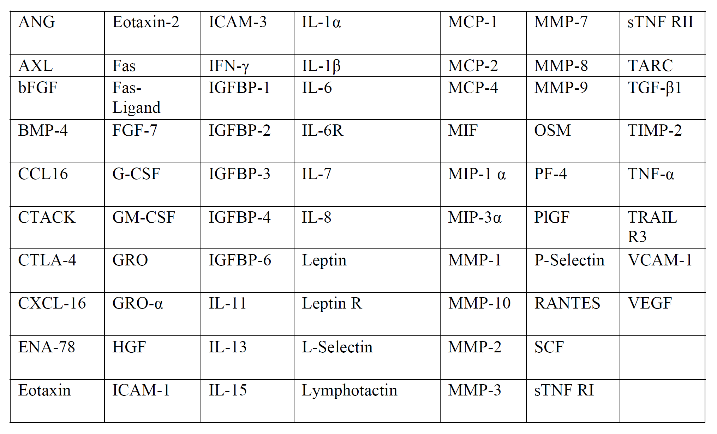

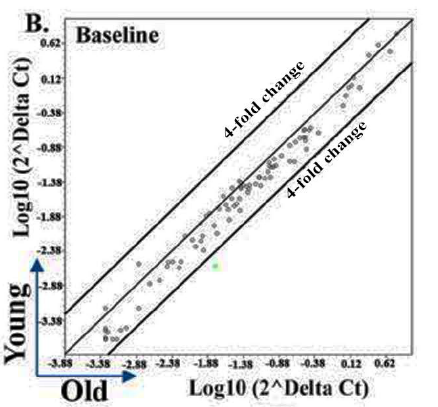

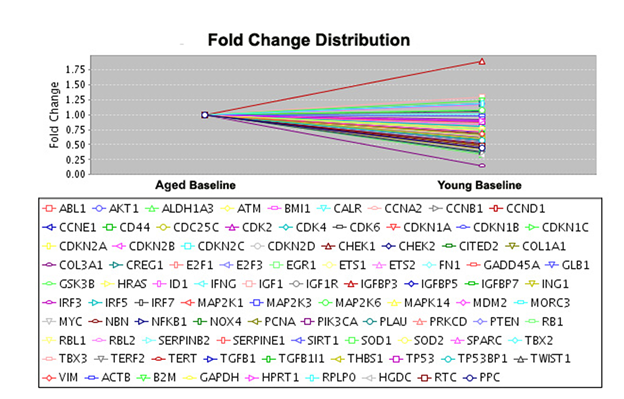

**C.**

**A.**

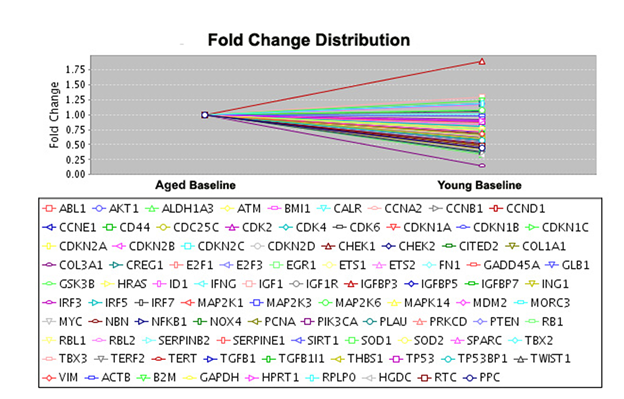

**D.**

**E.**

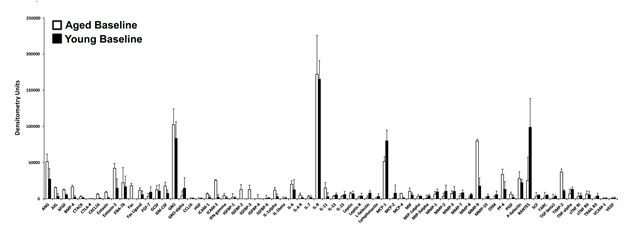

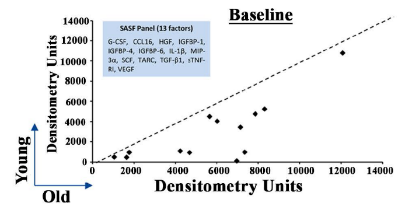

**F.**

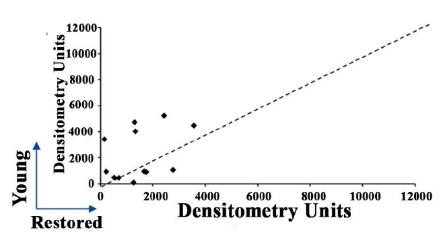

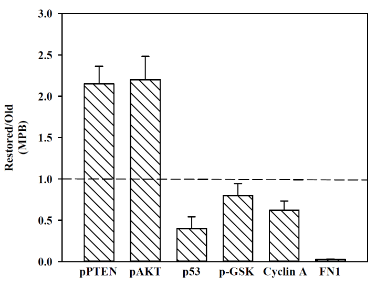

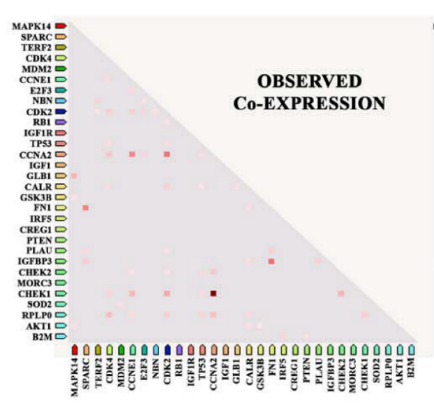

**G.**

**I.**

**H.**

**J.**

**J.**

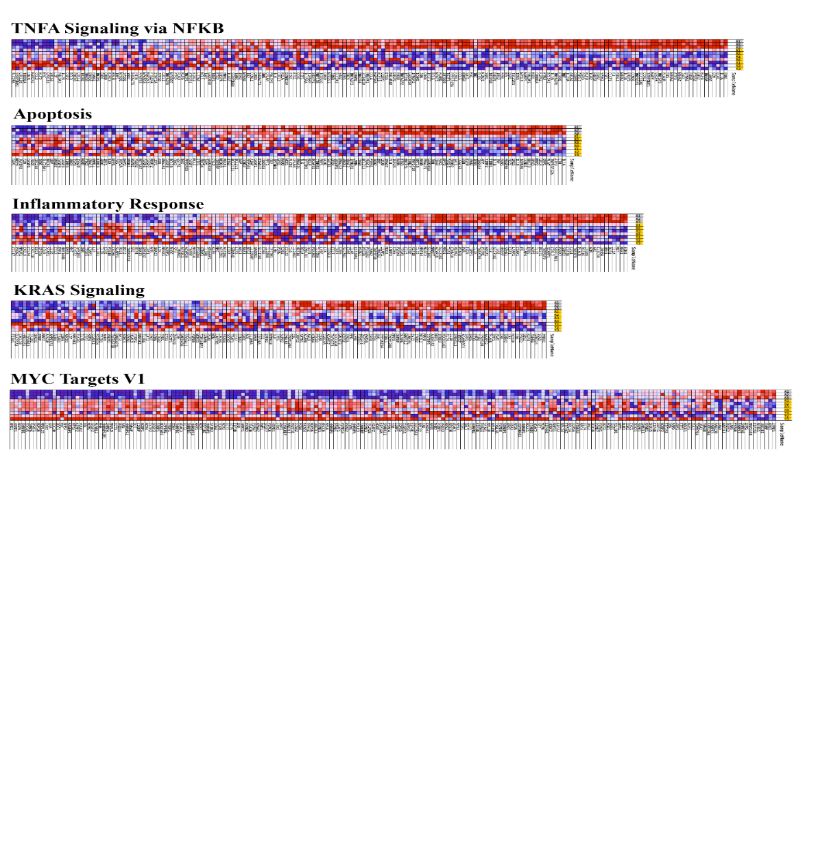
**Figure S2. (A)** The 84 age/senescence-related genes are those within the PCR arrays that were used to analyze baseline young and old MPBs, and restored MPBs. **(B)** The results of the analyses in `A’ were calculated as follows: ΔC_t_ between gene-of-interest and housekeeping genes and the values used to calculate fold change. Shown are the boundaries with 4-fold changes in gene expression. **(C)** Custom senescence-associated secretome (SASP) of 64 factors tested for released factors in aged and young bases line as well as restored samples. **(D)** Line plots representing 4 biological replicates, illustrate the fold change of proteins between aged and young baseline MPB. **(E.)** The denistometric results of analyses with media from heterochronic cultures at day 1. Densitometric units were calculated and plotted as a bar graph comparing expression of individual factors among aged and young. Results are presented as the mean ± SD, n=4. **(F)** The dot plot shows 13 differentially expressed proteins between young and old MPB. The dotted line *y=x* indicates no change between groups. **(G.)** Similar plots was made to determine similarities between young and restored using the method described in `F’. **(H.)** The plots shows co-expression of genes in the 30 genes in the boxed region of Figure 1I. The dots represent the association between genes. **(I.)** Normalized band densities for Western blots shown in Figure 1K (n=3, ±SD). **(J.)** The figure represents the full heat map in Figure 2I.

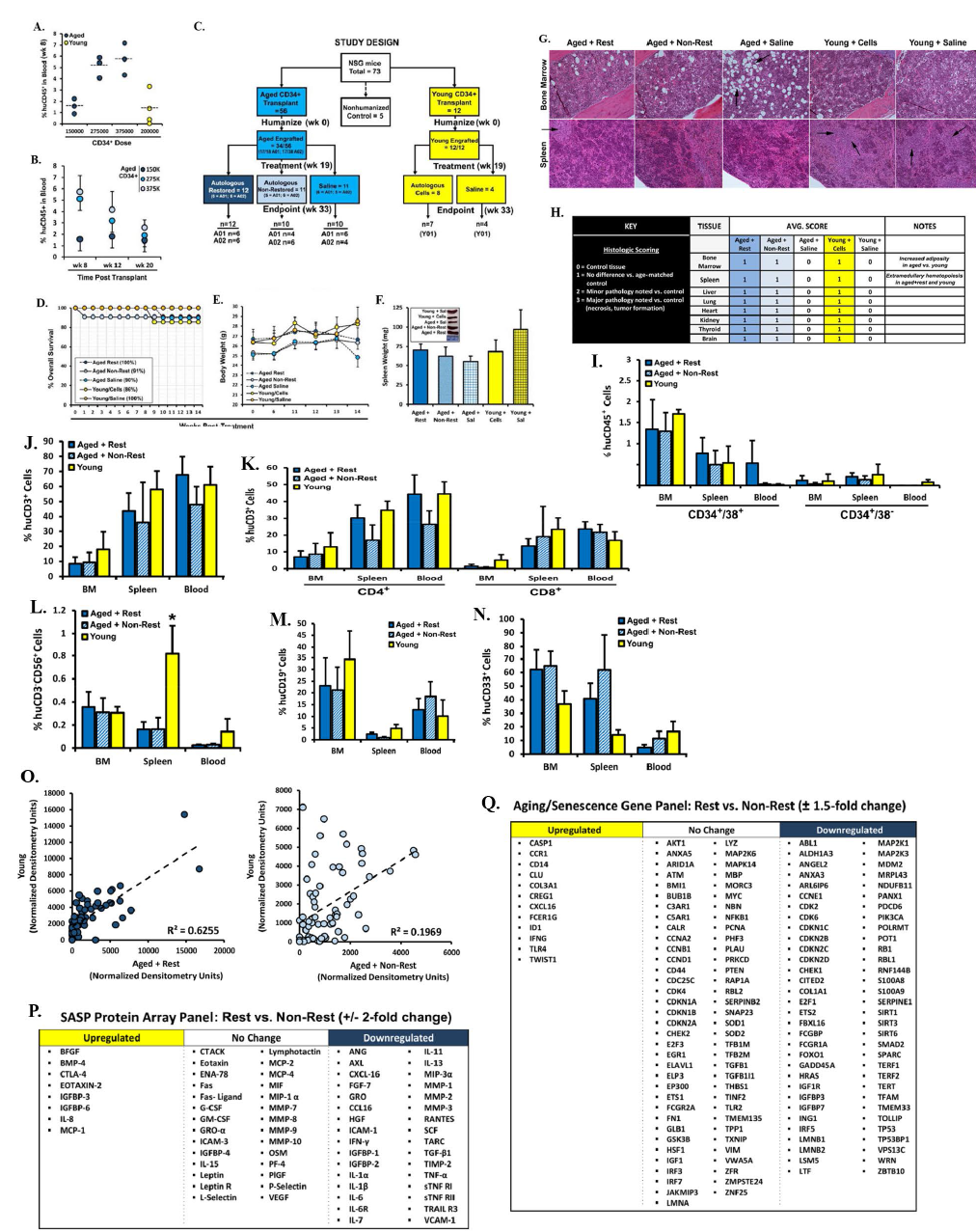

**Figure S3.** Procedural and safety monitoring of huNSG mice**:** (**A)** To identify the optimal dose, we transplanted different doses of aged CD34+ cells in NSG mice. The figure shown engraftment (human chimerism) at wk 8, based on human CD45+ cells in peripheral blood. (**B**) The transplant in `A’ was repeated with chimerism assessed up to 20 wks post-transplant, based on huCD45 in blood. (**C**) Study design for huNSG aging cell restoration study. A total of 68 irradiated mice were transplanted with aged (n=56) or young (n=12) CD34^+^ cells, with 34 of 56 mice successfully engrafted (based on human chimersim) with aged, and 12 of 12 mice successfully engrafted with young CD34^+^ cells. Chimerism cutoff for enrolling mice in the second transplant with cells from heterochronic and isochronic cultures was ≥1% huCD45^+^ cells in blood. Mice displaying 0.5%-1% chimerism were enrolled in the saline treatment arms, and mice displaying <0.5% chimerism were not enrolled in the study. The injected cells were from two different aged donors (A01 and A02), restored/heterochronic or unrestored/isochronic (**D**) Kaplan-Meier plot for huNSG survival. (**E**) Body weight at wk 14 following the 2^nd^ transplant. (**F**) Spleen weights are shown at the endpoint, with spleen images in legend inset. Results are presented as the mean ± SEM. **p*≤0.05 vs. control.

Histologic evaluation of tissues from huNSG treatment groups at the study end points: **(G**) Major organs and immune tissues were harvested. H&E staining of mouse femur (top panels) and spleen (bottom panels), 10X magnification. (**H**) All harvested tissues were examined by a pathologist for tissue necrosis and tumorigenesis. Treatment groups were compared to age-matched control tissue for pathological comparison. The black arrows at the top panel showed increased adipocytes femurs of mice given only aged cells whereas similar area of adipocytes was not noted in mice with a second transplant of restored cells. The arrows on the lower panels showed increased hematopoiesis in the spleen with restored or young cells.

Phenotype for human hematopoietic and immune cells in huNSG mice: Nucleated cells from blood, bone marrow (BM) and spleen were gated for human immune cells (huCD45^+^) and then analyzed for **(I)** hematopoietic stem (CD34^+^38^-^) and progenitor (CD34^+^38^+^) cells; (**J**) T-cells (CD3^+^); (**K**) T-helper (CD4^+^) and cytotoxic (CD8^+^) cells; (**L**) natural killer cells (CD3^-^56^+^); (**M**) B-cells (CD19^+^); and (**N**) myeloid cells (CD33^+^). Results are presented as the mean ± SEM.

Senescence- and aging-related gene and protein expression in huNSG treatment groups: (**O**) Scatterplots comparing senescence-associated secretory factor (SASF) expression in plasma of mice transplanted with either aged restored (left plot) or non-restored (right plot) cells compared to young. Values are normalized by background subtraction of SASF levels in non-humanized control NSG mice. Results are presented as mean densitometry units, with description of upregulated, downregulated and no change in expression SASFs listed in (**P**). (**Q**) List of aging- and senescence-related genes whose expression is upregulated, downregulated or no change in human cells isolated from huNSG BM. Classifications in **P** and **Q** are based on a 1.5-fold change cutoff.

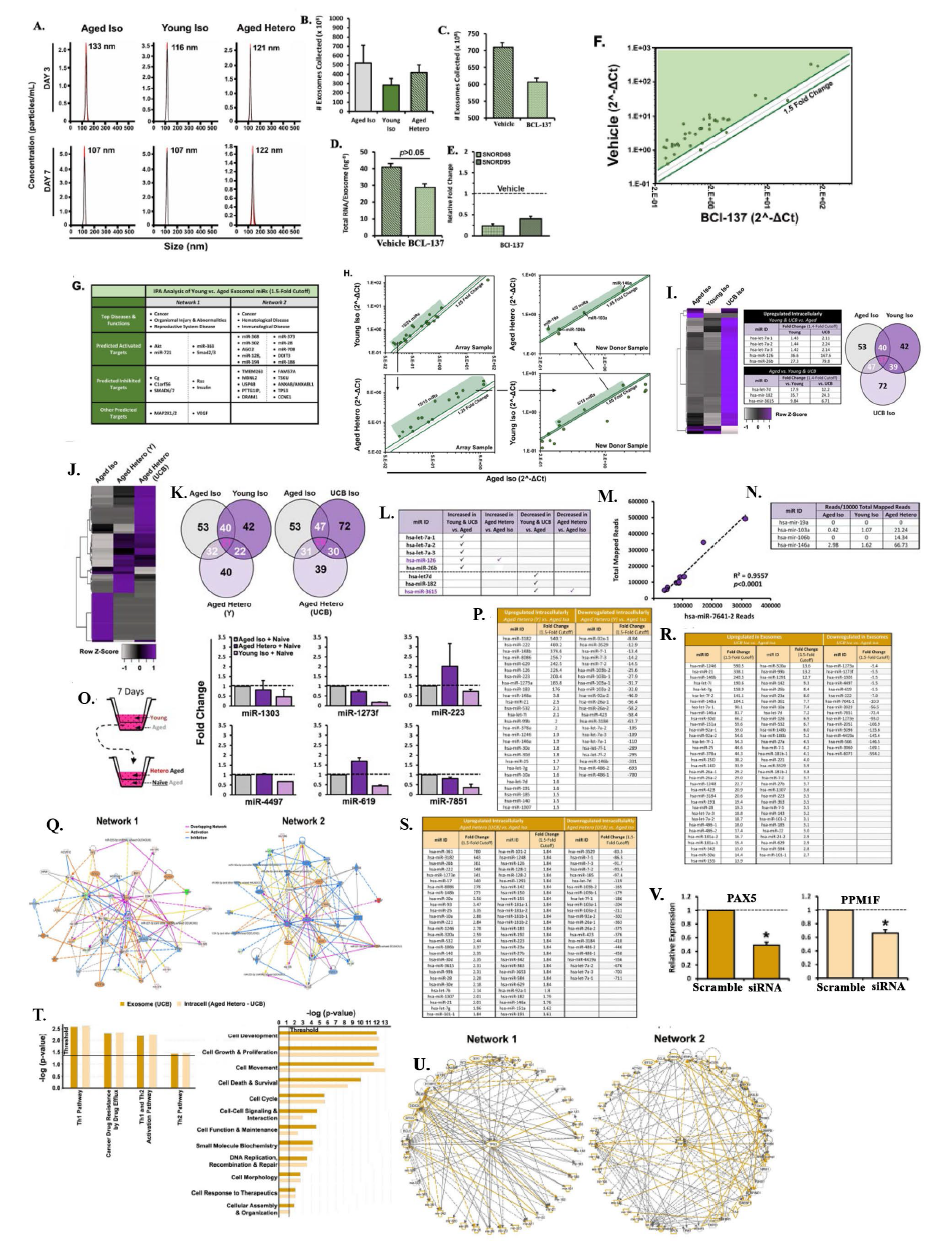
**Figure S4.** Exosomes and exosomal miRNAs in heterochronic (restored) and isochronic (unrestored) cultures (A-H): (**A**) Nanoparticle tracking analyses (NTA) of exosomes from day 3 and 7 heterochronic cultures. (**B**) Exosomes were isolated from 7-day heterochronic and isochronic control cultures on day 4 and day 7, and then pooled and quantified by nanoparticle tracking analysis (NTA), mean±SD, n=4. (**C**) Effect of AGO2 inhibitor (BCI-137) on exosome release. Exosomes were collected at days 4 and 7 from the media of heterochronic cultures, in the presence of BCI-137 or vehicle and then pooled for quantitation by NTA. The values are presented as the mean±SD, n=4. (**D**) Exosomes from `C’ were quantitated for total RNA. (**E**) Exosomes from `C’ were quantitated for small RNA. (**F**) Exosomes from `C’ were analyzed with miRNA array. The results are shown for enrichment of exosomal miRNAs in cultures without inhibitor vs. vehicle in a scatterplot, based on 1.5-fold cutoff. (**G**) Shown are the output of Ingenuity Pathway Analysis (IPA) using commonly expressed miRNAs with differential expression using 1.5-fold cutoff. The data were obtained from `F’ and the analyses compared young vs. aged isochronic cultures. (**H**) Validation of miFinder qPCR array by individual qPCR experiments in array (left panels) and fresh donor samples (right panels). Gating scheme depicts miRNAs that are upregulated in young isochronic and heterochronic vs. aged isochronic cultures. Results are depicted by scatterplot with 1.25-fold and 1.05-fold cutoffs in array and fresh donor samples, respectively. Array and individual qPCR studies were normalized to RNU6, SNORD68 and SNORD95 and presented as fold change, with a value of 1 representing control.

Characterizing the young intracellular miRNAs and ascribing a role for miRNAs in the mechanism of restoration (I-O): (**I**) Small RNA was purified from aged, young and UCB isochronic cultures for whole miRNA sequencing. All miRNAs exhibiting greater than 100 mappable reads were further analyzed. Differential RNA expression is denoted by heatmap, with miRNA exhibiting greater than 1.4-fold difference among aged vs. UCB and young samples. Outer area of the Venn diagrams depicts total number of intracellular miRNAs with greater than 100 mappable reads in age-matched isochronic samples. Overlapping areas represent common miRNA among samples. (**J, K**) Studies, similar to those in `I’, compared the miRNAs obtained from sequencing of aged isochronic and heterochronic (young-aged, UCB-aged) samples. (**L**) MiRNA showing differential expression in `**A’** were compared to miRNA showing increased or decreased expression in heterochronic (aged-young) vs. aged isochronic cultures in `J’ and `K’. The results are tabulated to illustrate candidate miRNAs whose expression patterns are coincident with aged cell restoration. (**M**) Scatterplot depicting linear correlation between exosomal miR-7641-2 expression and total mappable reads, n=10. **(N**) Expression of early exosomal candidate miRNAs from miFinder array studies in sequencing dataset. Results are shown for isochronic and heterochronic cultures as average reads per 10000 total mapped reads. (**O**) To evaluate whether candidate exosomal miRNAs can be propagated after aged cell restoration, aged and young cells from 7-day isochronic cultures or aged cells from heterochronic culture were harvested at day 7 and transferred to fresh transwell cultures with naïve aged cells for an additional 7 days. On the 3^rd^ (day 10) and 7^th^ (day 14) day of the propagation culture, exosomes were isolated and probed for candidate miRNA expression by qPCR. Results were normalized to miR-7641-2 expression and presented as fold change, with a value of 1 representing control (gray bars). Results are presented as the mean ± SEM, n=3, unless otherwise noted. **p*≤0.05 vs. control.

Identification of potential young exosomal miRNA targets in aged cells (P-V): (**P**) Up- and downregulated intracellular miRNAs comparing aged heterochronic (aged-young) vs. isochronic cultures with a 1.5-fold cutoff, and their (**Q**) predicted activation/inhibition networks after IPA. Up- and downregulated (**R**) exosomal miRNAs comparing UCB vs. aged isochronic and (**S**) intracellular miRNAs comparing aged heterochronic (aged-UCB) vs. isochronic cultures with a 1.5-fold cutoff. (**T**) Illustration of the top cellular functions (left graph) and canonical pathways (right graph) predicted by (**U**) these networks are shown. (**V**) Validation of siRNA knockdown of target candidates in cells from aged donors. Results were normalized to β-Actin expression and presented as fold change, with a value of 1 representing control (scrambled siRNA). Results are
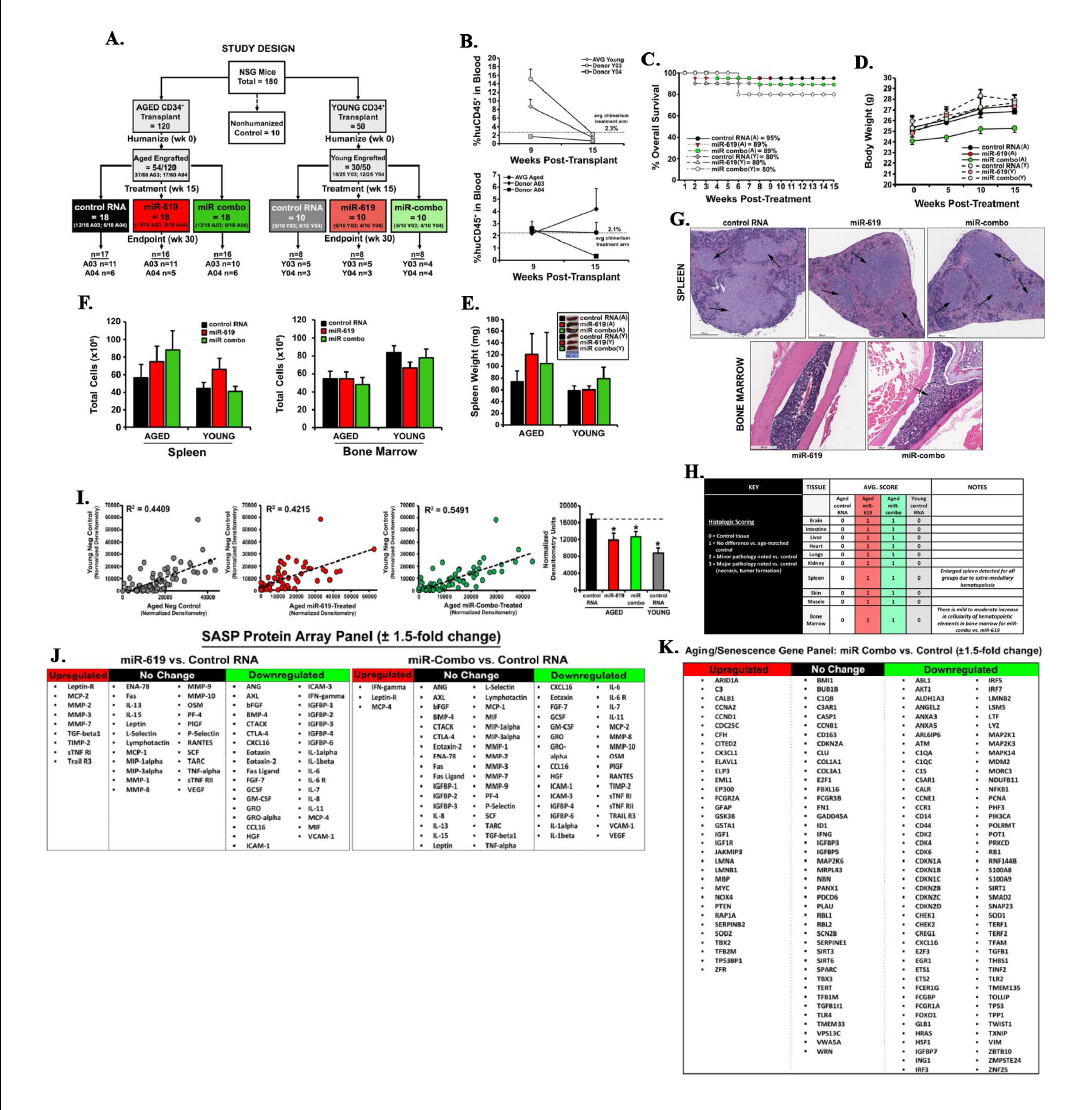
presented as the mean ± SEM, n=3, unless otherwise noted. **p*≤0.05 vs. control.

**Figure S5.** Procedural and safety monitoring of humanized mice from the cell-free restoration study. (**A**) Study design for the cell free restoration study in humanized mice. A total of 170 irradiated mice were transplanted with aged (n=120) or young (n=50) CD34^+^ cells, with 54 of 120 mice successfully engrafted with aged, and 30 of 50 mice successfully engrafted with young CD34^+^ cells. Chimerism cutoff for enrolling mice in the control and treatment arms was a minimum of 1% huCD45^+^ cells in blood. Mice displaying 0.5%-1% chimerism were enrolled in the saline treatment arms (not shown), and mice displaying <0.5% chimerism were not enrolled in the study at all. (**B**) Bleeds were performed on mice transplanted with aged (A03 or A04) and young donor (Y03 and Y04) CD34^+^ cells, and chimerism evaluated at 9- and 15-weeks post-transplant. Average chimerism of the aged (top graph) and young (bottom graph) donors enrolled in the study are shown. (**C**) Kaplan-Meier plot for huNSG overall survival post-treatment and (**D**) mouse body weights for the 15 weeks following the 2^nd^ transplant are shown. Percent survival is displayed in the legend inset. (**E**) Mouse spleen weights at study endpoint, with spleen images in legend inset. Total (**F**) spleen and (**G**) bone marrow cellularity at study endpoint are displayed. Results are presented as the mean ± SEM.

Histologic evaluation of tissues from huNSG treatment groups in expanded study: At study endpoint, major organs and immune tissues were harvested. (**G**) H&E staining of mouse spleen (top panels) and bone marrow (bottom panels), 4X magnification. (**H**) All harvested tissues were examined by a pathologist for tissue necrosis and tumorigenesis. Treatment groups were compared to age-matched control tissue for pathological comparison.

Senescence- and aging-related gene and protein expression in huNSG treatment groups from expanded study: (**I**) Scatterplots comparing senescence-associated secretory factor (SASF) expression in plasma of mice transplanted with either aged + negative control (gray dot plot), aged + miR-619 (red dot plot) or aged + miR-combo (green dot plot) cells compared to young control. Values are normalized by background subtraction of SASF levels in non-humanized control NSG mice. Results are presented as mean densitometry units, with average total SASF expression among each group also shown for comparison (far right bar graph). Enumeration of SASFs upregulated, downregulated or not changed for miR-619 vs. control (left table) or miR-combo vs. control (right table) is listed in `**J’**. (**K**) List of aging- and senescence-related genes whose expression is upregulated, downregulated or not changed in human cells isolated from huNSG BM in miR-combo treated mice vs. control. Classifications in `J’ and `K’ are based on a 1.5-fold change cutoff. Results are presented as the mean ± SEM. **p*≤0.05 vs. control.

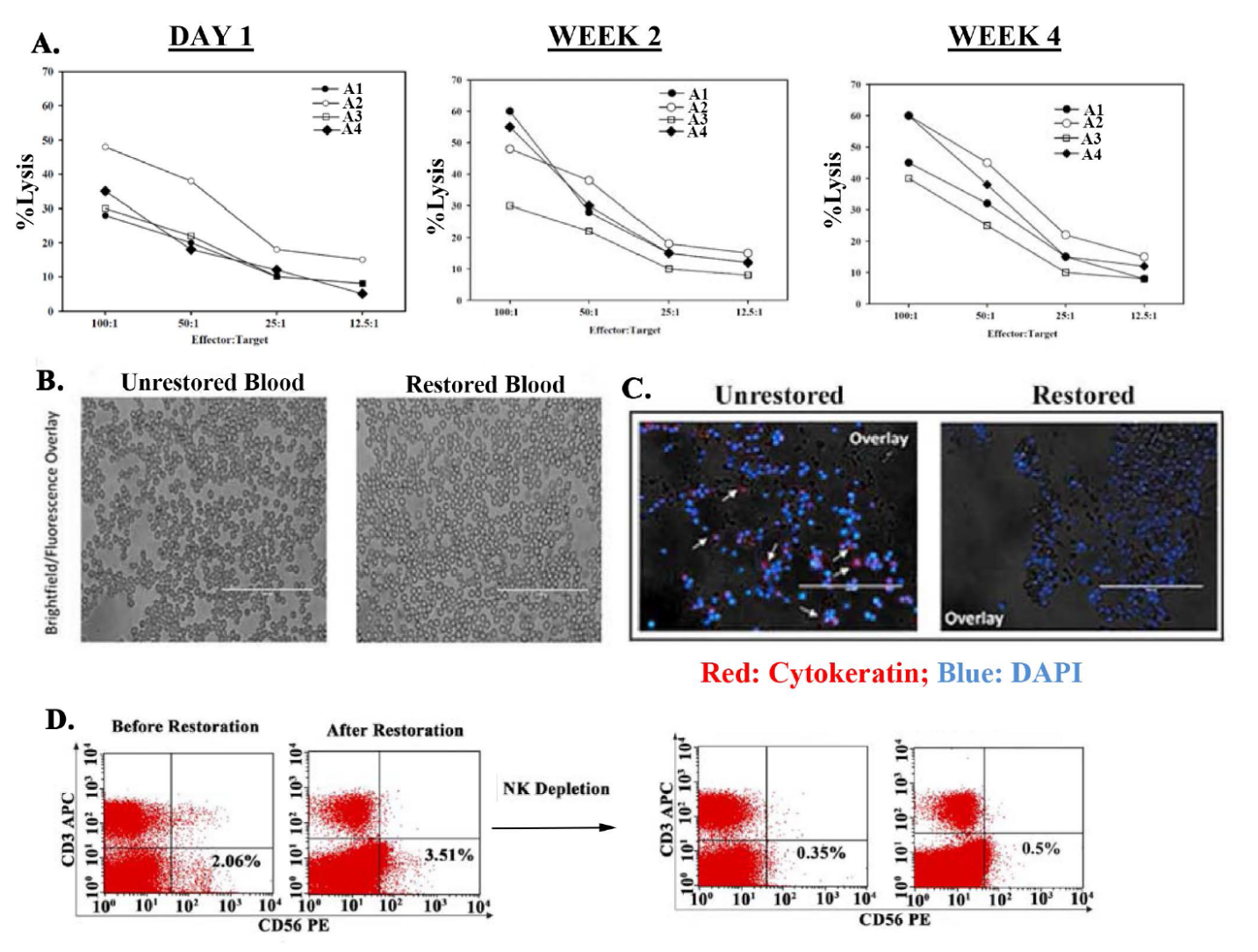
**Figure S6. A)** Timeline natural killer (NK) activity in restored old MPBs. NK activity was performed as described in Materials and Methods at day 1 and wks 2 and 4 following restoration in heterochronic cultures. The results are shown for four donors, each restored with three different young MPBs. **B)** Blood smears from mice, 12 months post-transplant, showed no detectable GFP^+^ cells, which served as a surrogate of CSCs. **C)** Femurs from mice at 12 months after transplantation with restored or unrestored MPBs. The femurs were dissected longitudinally and then washed with PBS. The cells close to the endosteal regions were scraped and placed onto microscope slides. The cells were fixed with 1% paraformaldehyde and then labeled with rabbit anti- human pan cytokeratin (1/1000) for human breast cancer cells and secondary labeling with PE-anti-mouse IgG. DAPI labeling for nuclear labeling of all cells. The slides were immediately imaged with the Evos fl2 auto imager. Representative images show the overlays of the labeled and bright field images at 100x magnification. **D)** Shown are representative results of restored and unrestored MPBs, with NK cells and NK-cell depletion.
